# Progressive Senescence Programs Induce Intrinsic Vulnerability to Aging-related Breast Cancer

**DOI:** 10.1101/2023.05.08.539818

**Authors:** Huiru Bai, Xiaoqin Liu, Meizhen Lin, Yuan Meng, Nan Li, Michael F. Clarke, Shang Cai

## Abstract

Cancer incidence escalates exponentially with advancing age; however, the underlying mechanism remains unclear. In this study, we built a chronological molecular clock at the single-cell transcription level with a mammary stem cell-enriched population to depict physiological aging dynamics in mice. We found that the mammary aging process was asynchronous and progressive, initiated by an early senescence program with elevated NF-kB and P53 signaling, succeeded by an entropic late senescence program with reduced NF-kB and P53 signaling and enhanced PI3K-Akt-mTOR, Wnt, Notch and pluripotent activity, vulnerable to cancer predisposition. The transition towards senescence program was governed by the master stem cell factor *Bcl11b*, loss of which accelerated mammary ageing with enhanced DMBA-induced tumor formation. We identified a drug TPCA-1 that can elevate *Bcl11b*, rejuvenate mammary cells and significantly reduce aging-related cancer incidence. Our findings established a molecular portrait of progressive mammary cell aging and elucidated the transcriptional regulatory network bridging mammary aging and cancer predisposition, which can be modulated to control cancer initiation; therefore, this study has potential implications for the management of cancer prevalence in the aged.

## INTRODUCTION

Aging has been recognized as the greatest risk factor for the vast majority of cancer types. As a significant extension of global lifespan, the burden of cancer incidence and cancer mortality have been rapidly increasing as major challenges to human health worldwide ^1,2^. Despite extensive advances in aging studies at the molecular, cellular and organismal levels, the exponential association between cancer occurrence and age ^3–5^ has persisted for years, and the underlying biology of this etiological phenotype remains largely unclear. Understanding how physiological aging programs impact carcinogenesis is therefore of particular interest and an imperative research direction for preventing cancer prevalence.

In contrast to the uncontrolled proliferation of cancer cells, in aging tissue, a senescence program is frequently activated, which suppresses cell proliferation, through the p53-p21 and p16^INK4a^–Rb pathways ^6,7^ and is accompanied by activated β-galactosidase ^8^, global alteration of H3K9me3 abundance ^9,10^, induction of senescence-associated secretory phenotype (SASP) factor activity ^11^ and disturbed immune ecosystem ^12^. In general, these pathways impair long-term stem cell self-renewal ability ^13^, induce chronic inflammation ^14^ and interfere with tissue homeostasis ^15^, ultimately leading to the acquisition of degenerative phenotypes. The prevailing idea to explain aging-caused cancer involves accumulation of mutations during aging, which promotes the transformation of cells into cancer cells ^16–19^; however, the degree to which this transformation drives cancer is not clear ^20^, largely because concordance between mutation rates and cancer profiles for aged individuals is lacking and because increased longevity is associated with lower cancer incidence ^3,21^. Alternatively, aging tissue may secrete a plethora of SASP factors, forming a fertile environment for neighboring cells to promote cancer initiation ^22^. Intriguingly, recent discoveries have indicated that the stemness program can be triggered by senescence during embryonic development ^23,24^, wound healing ^25–29^, and drug treatment in cancer ^30^. Whether this reprogramming underpins the physiological aging process and whether it contributes to cancer initiation remain unclear. Understanding chronological aging dynamics under physiological conditions and the underlying driving force mediating aging are crucial in establishing a biological connection between aging and cancer.

In this study, we addressed the potential connection between aging and cancer by building a chronological transcriptome map at the single-cell level with a stem cell-enriched mammary population in mice of various ages (from 2 to 29 months). We identified heterogeneous cell states in mice at each individual age with distinct senescence programs (early or late) vulnerable to breast cancer predisposition. In addition, we identified a master transcription factor, *Bcl11b*, that comprehensively suppressed both early and late senescence programs and found that loss of *Bcl11b* expression dramatically accelerated aging and tumor formation. Reversing the senescence program by TPCA-1 treatment efficiently reduced the cancer incidence and extended the cancer latency time. Our study established a molecular aging trajectory for mouse mammary cells and revealed an intrinsic molecular link between aging and cancer, which may shed light on preventive strategies against breast cancer occurrence in the future.

## RESULTS

### A Chronological Single-cell Transcriptome Analysis Reveals Asynchronous Dynamics of a Mammary Stem Cell-Enriched Population during Aging

Recently, single-cell studies have revealed hallmarks of aging in mammary gland cells at discrete ages, defined as young and old ^31,32^; however, the chronological aging process of mammary gland cells has not been well documented. Because initial oncogenic mutations occur in long-lived stem cells/progenitors ^33,34^, we were particularly interested in the molecular progression of aging in a mammary stem cell-enriched population. We therefore sorted mammary stem cell-enriched populations (CD49f^high^EpCAM^low^) ^35,36^, which included all the mammary stem cells identified to date ^37–40^, from mice of various biological ages spanning from 2 months to 29 months and performed single-cell RNA sequencing (RNA-seq) with 3’ UTR Smartseq ^41–43^ (Figure 1A). We extracted transcriptome information of 1981 cells from 25 mice (1-4 mice at each age) and detected 20629 genes. A subsequent t-distributed stochastic neighbor embedding (t-SNE) analysis revealed that the vast majority of the mammary cells obtained from mice of various ages were thoroughly intermingled (Figure 1B). These cells shared a similar transcriptome and uniformly expressed *Keratin14* and *Keratin5* (Figure 1C), suggesting that they share the same fate.

**Figure 1.**
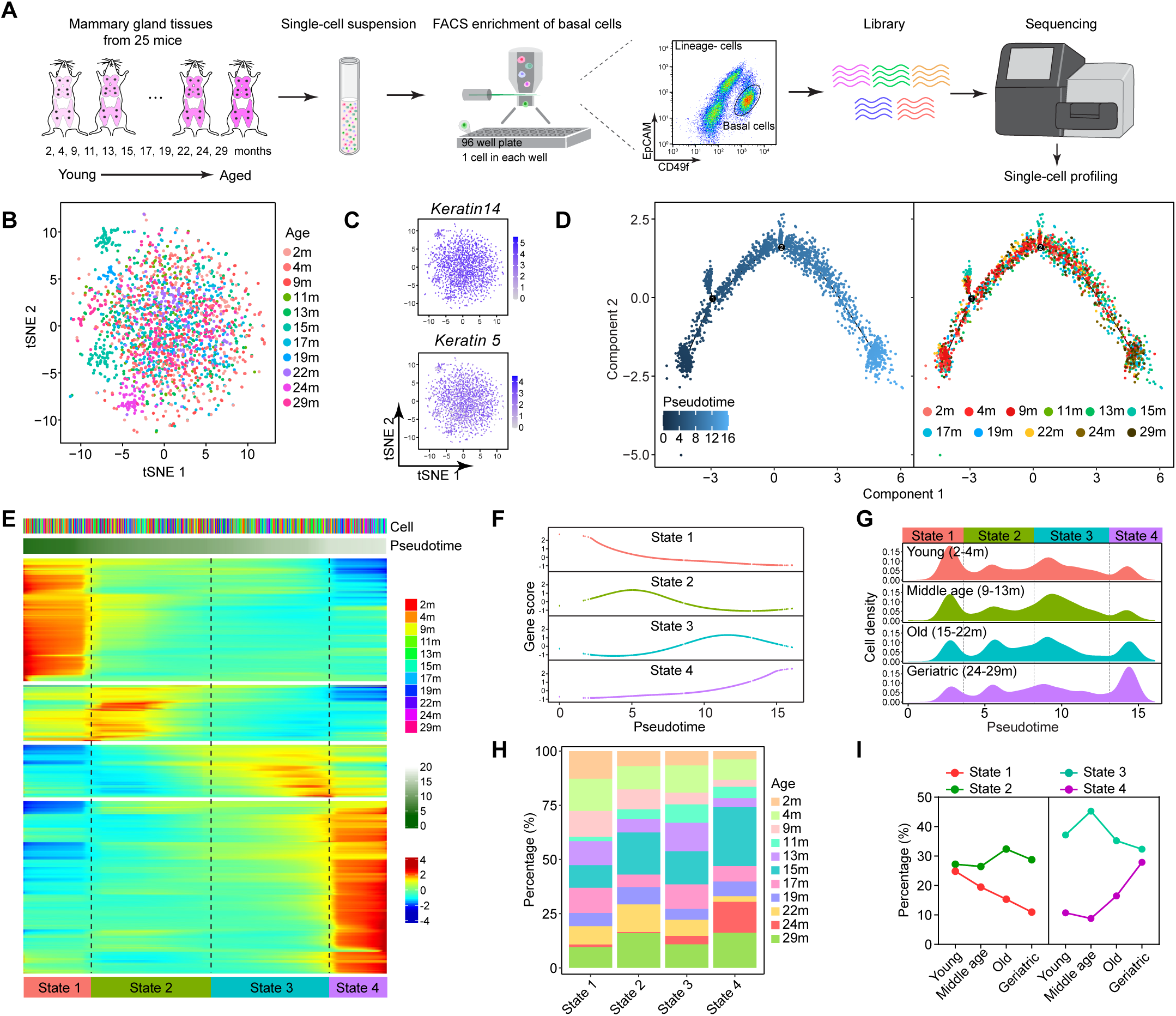
scRNAseq profiling of mammary stem cell enriched population at various chronological ages of mice. (A) Schematic diagram showing the pipeline of scRNA-seq of mouse mammary cells. Dissociated mammary cells from 2-month to 29-month old mice were sorted for CD49f^high^EpCAM^low^ mammary stem cell enriched population, followed by 3’UTR SMARTseq for library construction and subsequent sequencing. (B) t-SNE plot showing the clustering of mammary cells originating from 2m (n=2), 4m (n=3), 9m (n=2), 11m (n=2), 13m (n=2), 15m (n=4), 17m (n=2), 19m (n=2), 22m (n=2), 24m (n=1) and 29m (n=3) mice. (C) Basal cell specific genes (*Keratin 5* and *Keratin 14*) on t-SNEs showing a uniform expression pattern. (D) Pseudotemporal ordering analysis of single cell transcriptomes by Monocle 2 inferring the mammary ageing trajectory. Cell origins are labeled by distinct colors. (E) Heatmap visualization of the dynamic gene expression over the pseudotime. Cells were divided into four states based on the differentially expressed gene clusters. (F) Expression pattern of the signature gene clusters for each cell state along the pseudotime. (G) Cell density map showing the distribution of mammary cells from young (2m-4m), middle age (9m-13m), old (15m-22m) and geriatric (24m-29m) mice along with pseudotime. (H) Relative cell proportion of each mouse age in each cell state. (I) Relative cell abundance of the four mammary cell states in each age group.

To build a molecular clock and thus gauge dynamic transcriptomic changes with age, we performed a trajectory analysis with Monocle 2 and reconstructed a linear pseudotime ordering of mammary cells at different mouse ages. Remarkably, the mammary cells at different mouse ages clearly followed a chronological order, with the cells isolated from younger mice aligning with the early pseudotime stage and the cells isolated from older mice aligning with the later pseudotime stage (Figure 1D). This finding indicates that an age-related transcriptome program defines the intrinsic cell state. Indeed, when we clustered the differentially expressed genes on the basis of the pseudotime, the signature genes in the mammary cells were classified into four different states with distinct gene expression patterns (Figures 1E, 1F and S1B). Interestingly, the mammary cells of each individual mouse comprised all four-state cells, with their relative abundance being the only difference (Figures 1G, 1H and S1A). We then quantified the relative abundance of the cells in each cell state throughout the aging process and found that the number of State 1 cells kept decreasing with age and that of the State 4 cells continuously increased with age. The number of state 2 cells remained relatively constant, while that of State 3 cells temporarily increased and then decreased (Figures 1I and S1A, S1C-F). These data suggest that the mammary CD49^hi^EpCAM^low^ cells are heterogeneous with distinct cell states and that the aging phenotype may be manifested by the relative abundance of various cell states.

### Distinct Senescent Mammary Cell States Induce Progressive Aging Dynamics

To further characterize the biological identities of each individual cell state, we performed pathway analysis. We plotted the activity of each signaling pathway over pseudotime to visualize the chronological dynamics, and we identified six distinct dynamic patterns (Figure 2A). Pattern 1 pathways exhibited the highest activity in State 1, gradually declined throughout the entire time course to the last state. These pathways included ‘DNA replication’, ‘mismatch repair’, ‘oxidative phosphorylation’, ‘beta-alanine metabolism’ and ‘valine, leucine and isoleucine degradation’. The decreased activity of ‘DNA replication’ and ‘mismatch repair’ with increased pseudotime aligned with the notion that DNA mutations accumulate during aging ^44,45^. In addition, this finding indicated that State 1 cells are younger cells with higher DNA repair ability and metabolic activity. Consistent with Pattern 1, Pattern 2 pathways showed a transient increase in activity during the State1-2 transition, followed by a rapid decline. Pattern 2 exhibited only one activated pathway, ‘mitochondria ribosome’. The low pathway enrichment level may suggest low transcriptional activity and a relatively quiescent state. Indeed, when we plotted metabolism-related pathways, we found that State 2 cells exhibited relatively low metabolic lipid, amino acid and carbon activity but higher TGFbeta signaling pathway activity (Figure 2B) than State 1 cells. In addition, a cell cycle analysis ^46^ suggested that State 2 cells contain a relatively high portion of cells in the G0 phase compared to that of the other cells (Figures 2C and 2D). Given the low metabolism in State 2 cells, we labeled these cells quiescent young (q-Young) cells, and we labeled State 1 cells active young (a-Young) cells.

**Figure 2.**
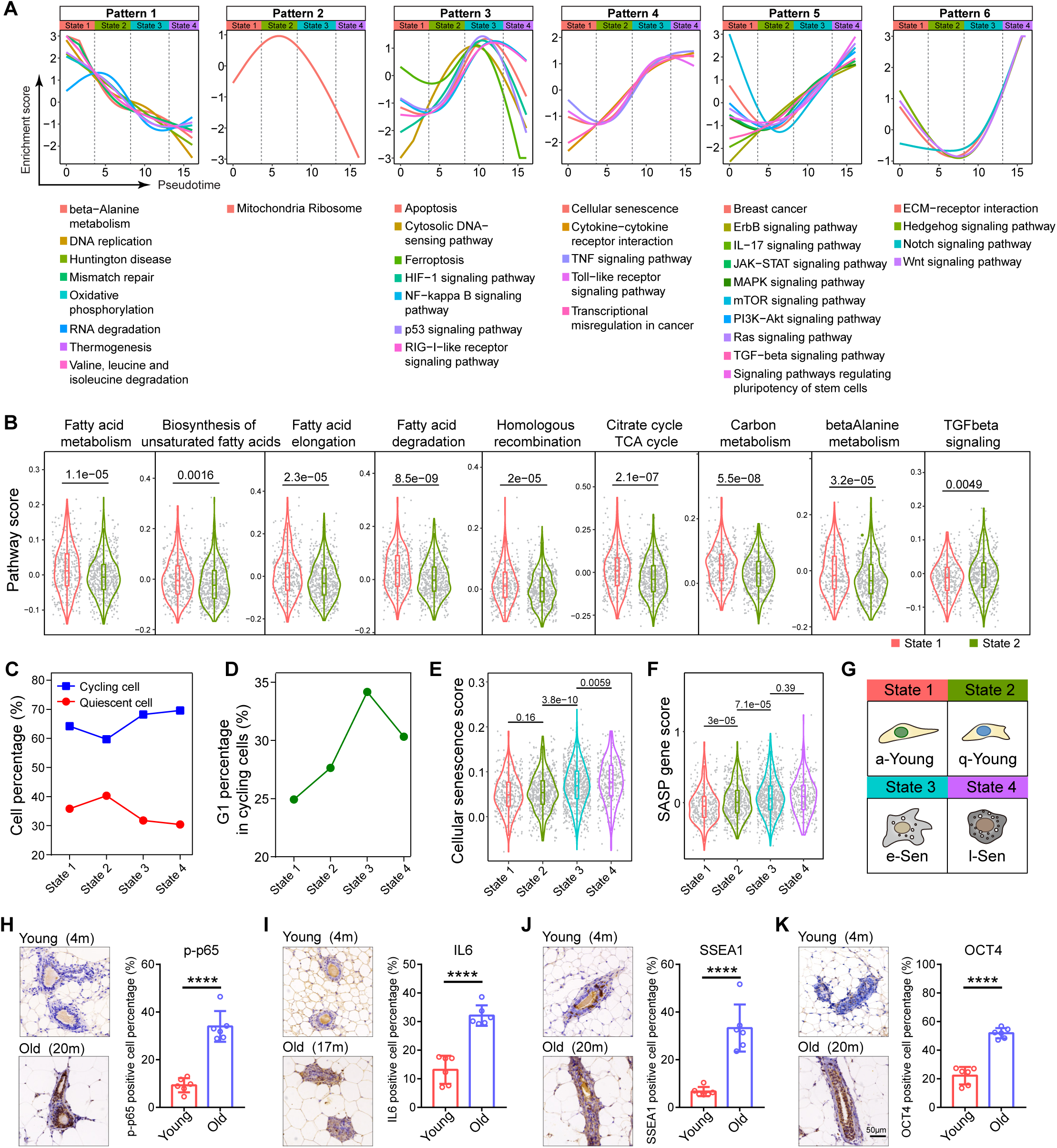
Characterization of distinct mammary cell states along the ageing trajectory. (A) Activity of each signaling pathway presented by enrichment score over the pseudotime. Note: six distinct dynamic patterns were identified indicating the identity of each cell state. (B) Differential pathway activities between state 1 and state 2 cells. The box plots show the interquartile range (box limits) and median (center blank line). Statistical analysis was performed using wilcoxon test. (C) Cell cycle analysis showing the relative fractions of cycling cells (blue) and quiescent cells (red) in each cell state. (D) Relative proportion of G1-phase cells in each cell state. (E-F) Senescence related pathway analysis showing the cellular senescence level (E) and SASP expression level (F) in each cell state. The box plots show the interquartile range (box limits) and median (center blank line). Statistical analysis was performed using wilcoxon test. (G) Schematic diagram of the four cell states designated as active young state (a-Young), quiescent young state (q-Young), early senescence (e-Sen) and late senescence (l-Sen). (H-K) Representative images of immunohistochemistry staining of p-p65 (H), IL6 (I), SSEA1 (J), OCT4 (K) in young and old mammary tissues.

Patterns 3-5 shared a similar increase in activity with nuanced differences. The activity of Pattern 3 pathways, including the ‘NF-kappaB signaling pathway’, ‘p53 signaling pathway’, ‘HIF1 signaling pathway’ and ‘ferroptosis’, peaked in State 3 and then quickly decreased in State 4 (Figure 2A). Notably, activation of the NF-kB and p53 pathways correlated with aging phenotypes ^47,48^. Pattern 4 pathways were activated in State 3 and then reached a plateau level that persisted into State 4. This pattern included ‘cellular senescence’, ‘cytokine–cytokine receptor interaction’ and ‘Toll-like receptor signaling pathway’ (Figure 2A). This finding suggests that both State 3 and State 4 cells had an activated senescence program. In accordance with this supposition, the expression of SASP program genes, a hallmark of aging cells, was elevated in both State 3 and State 4 cells (Figures 2E, 2F, S3A and S3B). Pattern 5 included pathways continuously increased activity in both States 3 and 4. These pathways included ‘IL-17 signaling’, ‘JAK-STAT signaling’, ‘mTOR signaling’, ‘PI3K-Akt signaling’, ‘MAPK signaling’, ‘Ras signaling’, ‘TGFbeta signaling’, ‘breast cancer’ and, most notably, ‘signaling pathways regulating pluripotency of stem cells’ (Figure 2A). The activation of these gene pathways suggests that State 4 cells, despite expressing the senescence program, had acquired stem cell traits and cell growth/survival programs, which may have predisposed them to a precancerous phenotype. We therefore named State 3 cells early senescence (e-Sen) cells, while State 4 cells were called late senescence (l-Sen) cells. Pattern 6 contained pathways activated later, between State 3 and State 4, indicating that they might be functionally involved in driving the transition from the e-Sen phenotype to the l-Sen phenotype (Figure 2G). These pathways included ‘Hedgehog signaling’, ‘Notch signaling’ and ‘Wnt signaling’ (Figure 2A). All of the six pathway patterns dynamics remained the same in different age groups (Figure S2), suggesting that the dynamics were intrinsic to the cell state irrespective of their biological age. Some of the signaling pathways were confirmed by immunohistochemical (IHC) staining (Figures 2H-2K and S3C-S3E).

These intriguing distinct senescence programs suggest that senescent cells are heterogeneous and that physiological aging progresses in a sophisticated manner, not through a binary switch. This unique physiological aging process is consistent with the in vitro aging dynamics induced by oncogenes ^49,50^, as well as the aberrant activation of aging and stem cell programs during embryogenesis ^23,24^, wound healing ^26,28^ and cancer drug treatment ^30^, indicating a pervasive underlying mechanism.

### Breast Cancer Initiation is Associated with l-Sen Program

l- Sen cells exhibited aberrantly activated cancer- and stem cell-related programs, and have reduced P53 activity and enhanced PI3K-Akt activity. Considering that P53 and PIK3CA are the two most prominent mutation genes in breast cancer ^51^, we speculate that l-Sen cells have increased their vulnerability toward cancer transformation. This prompted us to ask, do these programs predispose cells to a precancer state? We therefore analyzed the paired human breast samples (tumor and tumor adjacent normal tissue) in TCGA database for pathway activity and transcription factor activity (Figure 3A). Interestingly, compared to the adjacent normal tissue, breast tumors are significantly elevated for various senescence related pathways, and the l-Sen program related pathways, including Notch signaling, Wnt signaling (TCF7L1, LEF1), Hedgehog signaling (GLI1, GLI2), and pluripotency related factors (MYC, SOX2 and KLF4), while e-Sen specific NFkB pathway is tuned down (Figures 3B-3D and S4A). This suggests that breast tumor tissues are closely associated with l-Sen signature.

**Figure 3.**
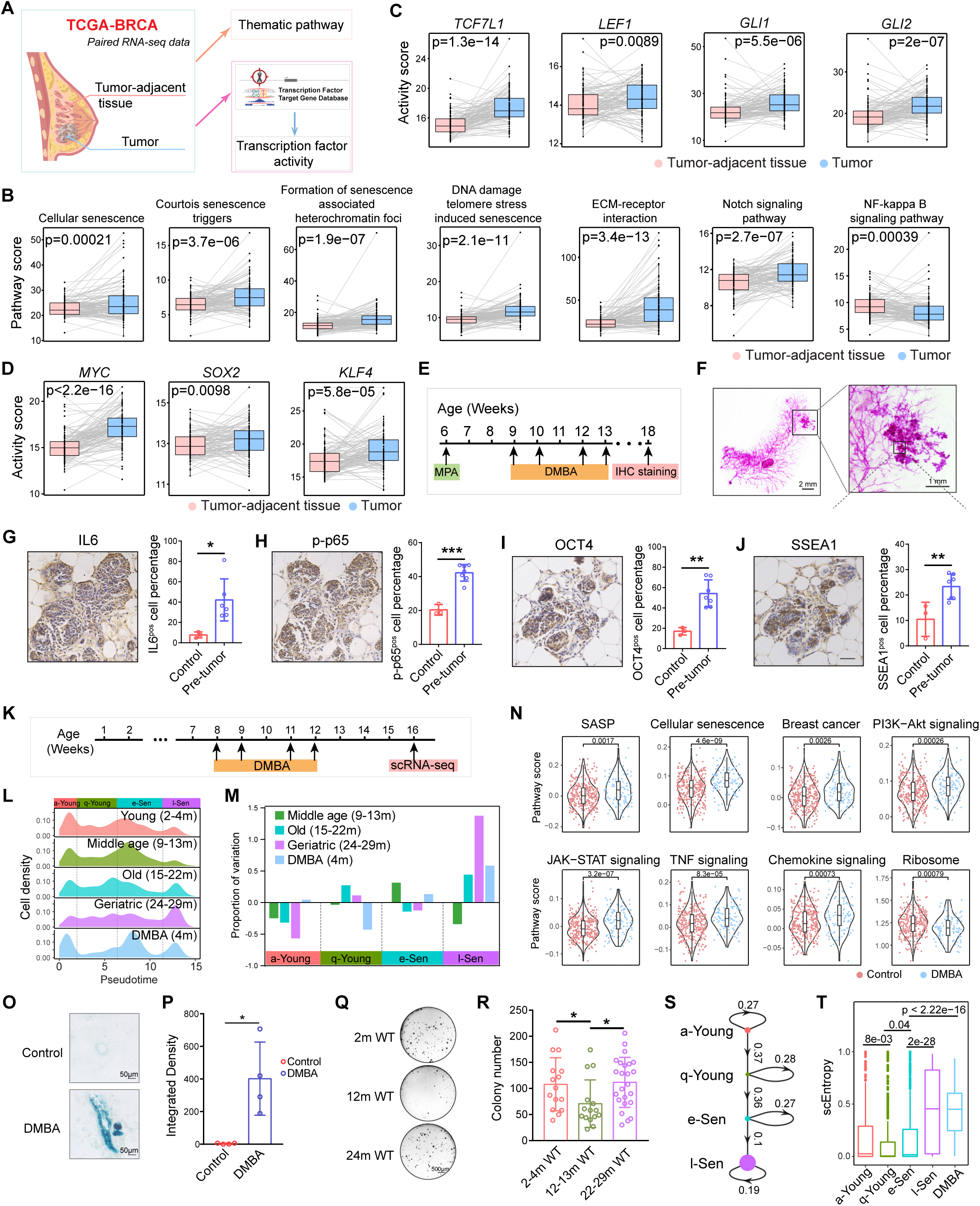
Senescent cells are vulnerable for cancer predisposition. (A) Diagram showing the workflow of the analysis for thematic pathway score and transcription factor activity score in human mammary gland tumor and tumor-adjacent tissue from TCGA database. (B) Plots showing the senescence related pathway score was higher in tumor tissue. Statistical analysis was performed using two-tailed paired t-tests. (C-D) Wnt and Notch pathway (C), Stemness (D) related transcription factor activity was upregulated in tumor tissue. Statistical analysis was performed using two-tailed paired t-tests. (E) Schematic diagram showing the DMBA induced cancer assay for 6-week-old WT mice. Mice were treated with MPA (50 mg, 90-day-release) plus DMBA (200μL, 5 mg/mL) at the indicated time in the schematic diagram. (F) Representative wholemount staining images for DMBA-induced tumors in mammary gland. (G-J) Representative IL6 (G), p-p65 (H), OCT4 (I), SSEA1 (J) IHC staining images for DMBA-induced tumors in mammary gland. (K) Diagram showing that 8-week-old mice were treated with DMBA (200μL, 5 mg/mL) and analyzed by scRNA-seq. (L) Density map of mammary cell states in each age group and DMBA treated group. (M) Changes of the relative abundance of cell states in different age groups compared to the Young (2-4m) group. (N) Senescence related pathway score in CD49f^high^EpCAM^low^ cells of DMBA treated mammary gland. (O) Representative β-gal staining images of mammary gland from control mice (4 month) and DMBA treated mice (4 month) showing senescent cells. Scale bar, 50μm. (P) Quantification of integrated density. Statistical analysis was performed using two-tailed unpaired t-tests; data are presented as mean ± SD; *P<0.05. (Q) Representative images of colony formation assay of CD49f^high^EpCAM^low^ mammary cells from 2m, 12m and 24m old WT mammary gland. 3000 cells/well were seeded and cultured for 7 days. Scale bar, 500μm. (R) Quantification of colony formation ability of CD49f^high^EpCAM^low^ mammary cells in 2-4m WT (n=5), 12-13m WT (n=5), 22-29m WT (n=8) mammary gland. Statistical analysis was performed using two-tailed unpaired t-tests; data are presented as mean ± SD; *P<0.05. (S) A linear lineage trajectory shows the transition probabilities for the four cell states with the node size corresponding to the signaling entropy. (T) scEntropy analysis of the four cell states. The box plots show the interquartile range (box limits) and median (center blank line). Statistical analysis was performed using wilcoxon test.

To ask whether the l-Sen program is turned on at the initiation stage of breast tumor, we used a dimethylbenz(a)anthracene (DMBA)-mediated breast cancer development model ^52,53^, which was shown to predominantly trigger breast tumors ^54^, and analyzed tumor initiation foci at very early stage (Figure 3E). Consistent with human breast tumors, the senescence related signals and pluripotency related signals were all upregulated at the onset of tumor formation (Figures 3F-3J). Meanwhile, we found that DMBA, besides its function causing genetic mutations, also triggered mammary cell senescence with a prominent l-Sen cell expansion and senescence related pathway activation, suggesting the tumor initiation process is accompanied with l-Sen program activation (Figures 3K-3P). Consistent with this idea, we found that mammary cells colony formation ability significantly decreased from young to old mice, but recovered in the geriatric mice where they have significant expansion of l-Sen population (Figures 3Q and 3R).

To further characterize the kinetics of the physiological aging process, we employed a single-cell signaling entropy algorithm ^55^ to profile the dynamics of cellular entropy (Figures 3S and 3T). The cellular entropy of a-Young, q-Young and e-Sen cells remained at a relatively low level, with a slight decrease from the a-Young to the q-Young cells (p<0.01) and an increase from the q-Young to the e-Sen cells (p<0.05). Remarkably, the entropy of the l-Sen cells was strikingly elevated compared with that of the e-Sen cells (p<0.001), indicating a drastic systemic disorder and a potential chromatin reorganization ^56^. The increase in entropy suggests that e-Sen cells transition to the l-Sen state in a passive spontaneous manner in the absence of extrinsic inputs. Therefore, we speculated that mammary aging might be initiated and determined during the early q-Young-to-e-Sen transition, which is crucial for subsequent l-Sen state commitment.

### The q-Young-to-e-Sen Cell Transition Is Mediated by the Mammary Stem Cell Factor *Bcl11b*

To determine the factors that drive the transition from the q-Young to the e-Sen state, we constructed a limited gene regulation network using the chromatin immunoprecipitation followed by sequencing (ChIP-seq) ENCODE and ChEA databases, along with the text curated database TRRUST of transcriptional regulatory networks, and we superimposed our network onto the expression matrix of the four aging-cell states to infer the key transcription factors in each cell state (Figure 4A). The highest fidelity factors supported by multiple databases were selected (Figure 4B).

**Figure 4.**
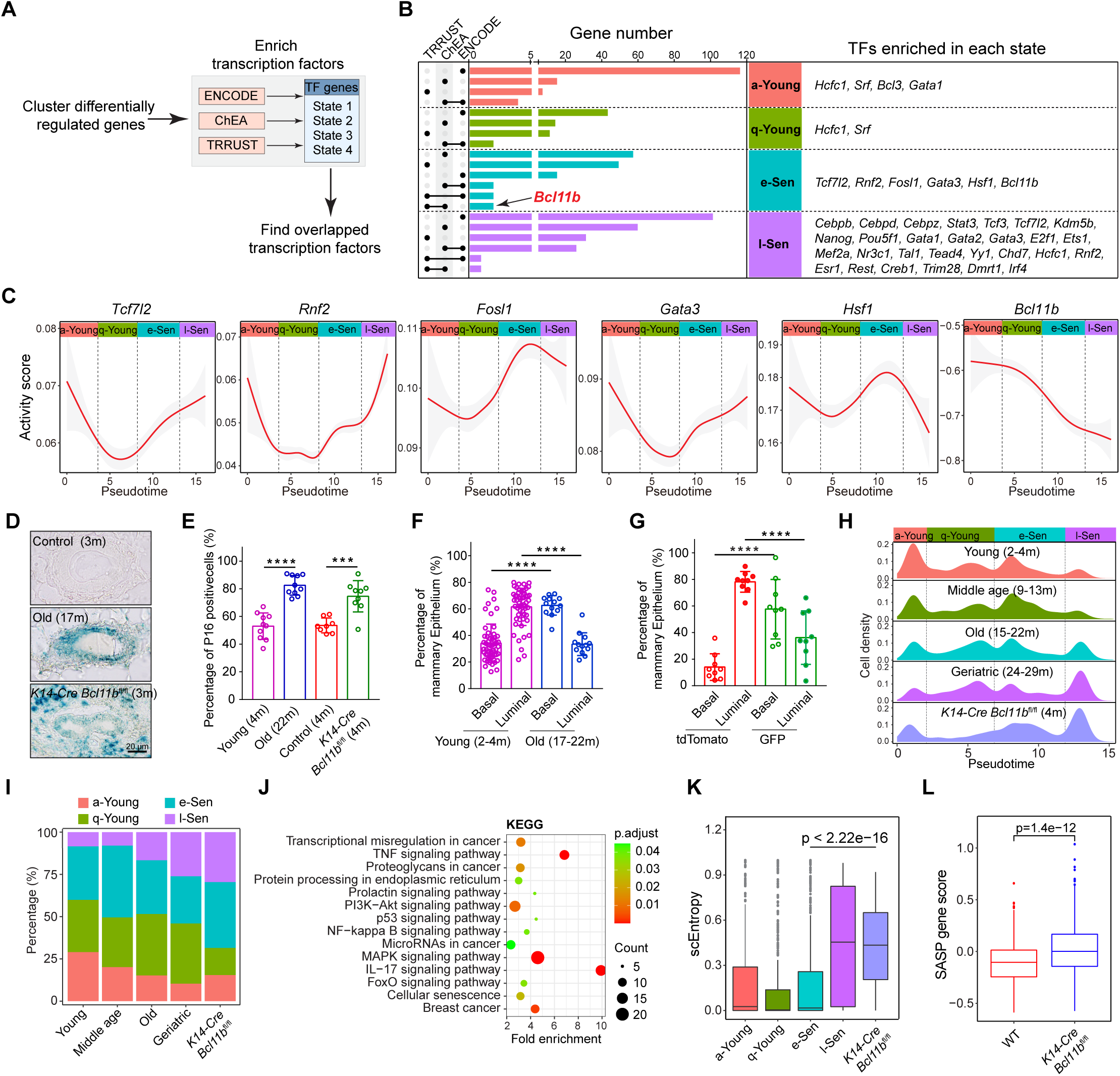
q-Young to e-Sen transition is functionally mediated by transcription factor *Bcl11b*. (A) Diagram showing the workflow of the transcription factors enrichment analysis for each cell state using the ENCODE, ChEA, and TRRUST database. Reliable candidates of transcription factors regulating each state were supported by at least two databases. (B) Table showing enriched transcription factors based on differentially expressed genes in each cell state. Black lines indicate TFs enrichment in two databases. (C) Activity score of transcription factors enriched in e-Sen state presented by target gene index over the pseudotime. We used the differentially expressed genes in each cell state, and inferred the transcription factors with the CHIP-seq target database. If the targets of certain TF factor are enriched in specific cell state, the TF factor will be popped out as a signature factor. (D) Representative β-gal staining images of mammary gland from young (3 month), aged (17 month) and *K14*-*Cre Bcl11b*^fl/fl^ (3 month) mice showing senescent cells. Scale bar, 20μm. (E) Percentage of p16^Ink4a^ (scale bar, 20μm) positive cells in mammary epithelial cells from young (4 month), old (22 month), control *K14*-*Cre Bcl11b*^wt/wt^ (4 month) and *K14*-*Cre Bcl11b*^fl/fl^ (4 month) mice. Statistical analysis was performed using two-tailed unpaired t-tests; data are presented as mean ± SD; ***P<0.001, ****P<0.0001. (F) Relative basal luminal proportion of mammary gland epithelia in young (n=54) and old (n=13) mice. Statistical significance was determined by two-tailed unpaired t-tests; data are presented as mean ± SD; ****P<0.0001. (G) Quantification of relative basal/luminal proportion in mammary epithelia in *K14*-*Cre Bcl11b*^fl/fl^ mTmG reporter mice (n=9). GFP+ cells were regarded as *Bcl11b* KO cells while tdTomato+ cells were regarded as WT cell control in the same gland. Statistical significance was determined by two-tailed unpaired t-tests; data are presented as mean ± SD; ****P<0.0001. (H-I) Density map (H) and percentage (I) of mammary cell states in each age group and *K14*-*Cre Bcl11b*^fl/fl^ group. (J) Kyoto Encyclopedia of Genes and Genomes (KEGG) pathway analysis showing pathways significantly enriched in *K14*-*Cre Bcl11b*^fl/fl^ CD49f^high^EpCAM^low^ cells. (K) Boxplots showing the scEntropy score of *K14*-*Cre Bcl11b*^fl/fl^ CD49f^high^EpCAM^low^ cells increased to a level similar to l-Sen cells. The interquartile (box limits) and median (center blank line). (L) Comparison of SASP gene score in WT and *K14*-*Cre Bcl11b*^fl/fl^ CD49f^high^EpCAM^low^ cells. The box plots show the interquartile (box limits) and median (center blank line). Statistical analysis was performed using wilcoxon test.

With a transcriptional inference algorithm, we pinpointed 38 state-specific fate determinants in total (Figure 4B), of which the majority were found in l-Sen state cells (Figure S4B). Interestingly, the factors in young state cells, such as *Hcfc1* and *Srf*, have been previously associated with longevity ^57,58^. In the l-Sen state cells, we found that factors such as *Cebpb*, *Cebpd* and *Cebpz* were associated with SASP secretion ^49,59^; *Stat3*, which is the downstream effector in the Jak-stat pathway; *Tcf3* and *Tcf7l2*, which are key effectors in the Wnt signaling pathway; and *Kdm5b*, which is a histone demethylase that contributes to Rb-mediated cellular senescence ^60^. Most interestingly, we observed Nanog and Pou5f1 which were associated with pluripotent stem cells. These factors were consistent with the pathway analysis, reinforcing the idea that in l-Sen state cells, the stem cell program is aberrantly activated.

In identifying the factors critical for the initiation of the e-Sen cell state, we concentrated our attention on e-Sen transcription factors. The activity plot showed that *Tcf7l2*, *Rnf2*, *Fosl1*, *Gata3* and *Hsf1* were all increasing their activity during the q-Young-to-e-Sen transition, with the exception that *Bcl11b* sharply decreased its activity (Figure 4C). *Bcl11b* is a previously identified modulator of mammary stem cell self-renewal and quiescence, the loss of which results in stem cell exhaustion ^38^. As *Bcl11b* is a predominant transcriptional repressor ^61–63^, a decreased *Bcl11b* activity is a reflection of the activation of *Bcl11b*-repressed targets. This finding suggests that the initiation of early senescence may be mediated by the loss of *Bcl11b* function.

### Accelerated Mammary Aging Phenotypes was Triggered by Loss of *Bcl11b*

We then wondered whether *Bcl11b* is a key factor mediating the mammary senescence switch. To determine the mechanism of *Bcl11b* function, *Bcl11b* expression was knocked out in mammary glands, and the *Bcl11b*-knockout (KO) mammary tissues exhibited reduced ductal width (Figures S5D-S5H), enhanced β-galactosidase activity (Figure 4D), and elevated p16^INK4a^ (Figures 4E and S5A), which are typical markers of senescent cells ^7,8^. In multiple aging organs, the stem cell population has been frequently shown to expand with declining functionality ^64–67^. Specific to aging mammary glands, a phenotype of an expanded basal population has been reported to be acquired with age ^68^, which is consistent with our observation of the mammary gland as the mice aged (Figures 4F and S5B). We analyzed *Bcl11b-*KO cells in a *Krt14*-*cre Bcl11b*^fl/fl^ mT/mG mouse, in which the green fluorescent cells represented *Bcl11b*-KO cells, and the red fluorescent cells were the *Bcl11b* WT control cells (Figure S5C). We found that, compared with the control cells in the same mouse, the *Bcl11b*-KO cells exhibited significant basal expansion at a young age (Figure 4G). Given that our previous data showed that knocking out *Bcl11b* expression significantly reduced mammary stem cell self-renewal ability ^38^, consistent with aging mammary stem cells (Figures S5I and S5J), these apparently expanded basal cells may undergo functional decline. Overall, these data collectively suggest that loss of *Bcl11b* function triggered an accelerated aging process.

Ageing has usually been defined on the basis of certain biomarkers or functional assays; however, these methods are of limited value when aging cells are heterogeneous. Because we built a molecular clock of mammary cell aging, we tried to use a chronological map to gauge the aging grades of the *Bcl11b*-KO cells. When we included the *Bcl11b*-KO cells in the aging pseudotime analysis and reconstructed the trajectory, we found that the vast majority of the *Bcl11b*-KO cells spontaneously accumulated at the later stage and coclustered with l-Sen cells (Figure S6), with a notably diminished q-Young peak (Figure 4H). Hence, quantification of each aging stage revealed that the number of a-Young and q-Young cells was drastically reduced compared with that of the age-matched wild-type (WT) cells, while the number of e-Sen and l-Sen cells was profoundly increased (Figures 4H, 4I and S6F). This finding suggested that *Bcl11b*-KO underwent substantially accelerated aging progression, and cells rapidly entered a state very similar to that of senescent cells.

To determine which aging-related molecular pathways were altered by knocking out *Bcl11b*, we performed a pathway analysis (Figures 4J and S7). The Kyoto Encyclopedia of Genes and Genomes (KEGG) analysis showed that the activity levels of the typical aging-related pathways, including ‘MAPK signaling’, ‘PI3K-Akt signaling’, ‘IL17 signaling’, ‘cellular senescence’, ‘NF-kB signaling’, and ‘p53 signaling’, were all significantly upregulated in the *Bcl11b*-KO cells, indicating a systemic aging state. In addition, the senescence-related SASP program was also elevated in the *Bcl11b*-KO cells, which was accompanied by increased cellular entropy (Figures 4K and 4L). Overall, these data demonstrate that *Bcl11b* may be a molecular switch regulating the q-Young-to-e-Sen transition, ultimately promoting the l-Sen chaos.

### Ageing Mammary Cells Induced by Bcl11b KO are Susceptible to Cancer Transformation

Next, we try to address whether the accelerated mammary ageing is intrinsically coupled with cancer in the DMBA treatment based cancer initiation model. In this tumor model, we found that the DMBA induced tumor cells predominantly originated from Krt14+ basal cells (Figures S8A-S8C), and the mammary cells showed prominent age-related susceptibility to chemical induced cancer transformation (Figures 5A-5C). We therefore determined to address whether the senescent mammary cells caused by *Bcl11b* loss are vulnerable to cancer transformation with the DMBA-induced breast tumor formation assay for WT and *Bcl11b*-KO cells. We tried to prevent microenvironmental factors from influencing the results by transplanting control and *Bcl11b*-KO cells into recipient mice at a saturated dose and then performed DMBA treatment after full mammary tree formation (Figure 5D). Intriguingly, the mice that received the *Bcl11b*-KO cells acquired tumors much earlier and faster than the control recipient mice, ultimately exhibiting a much higher cancer incidence with age (Figures 5E and 5F). We further characterized the WT and *Bcl11b*-KO tumors by scRNA-seq to address the possibility that the tumors originated from cells neighboring senescent cells (Figure S8D). We reasoned that if neighboring cells give rise to tumors, then WT and *Bcl11b* tumor cells likely have a similar phenotype. Interestingly, a gene set enrichment assay (GSEA) showed that WT tumor cells were significantly enriched with a-Young signature genes, while *Bcl11b*-KO tumor cells were enriched with l-Sen genes (Figure 5G). In addition, the *Bcl11b*-KO tumor cells themselves exhibited high levels of ageing cytokine IL6 and high NF-kB activity (Figures 5H and 5I). These data support the idea that the l-Sen cells induced by knocking out *Bcl11b* expression were intrinsically susceptible to transformation into cancer cells.

**Figure 5.**
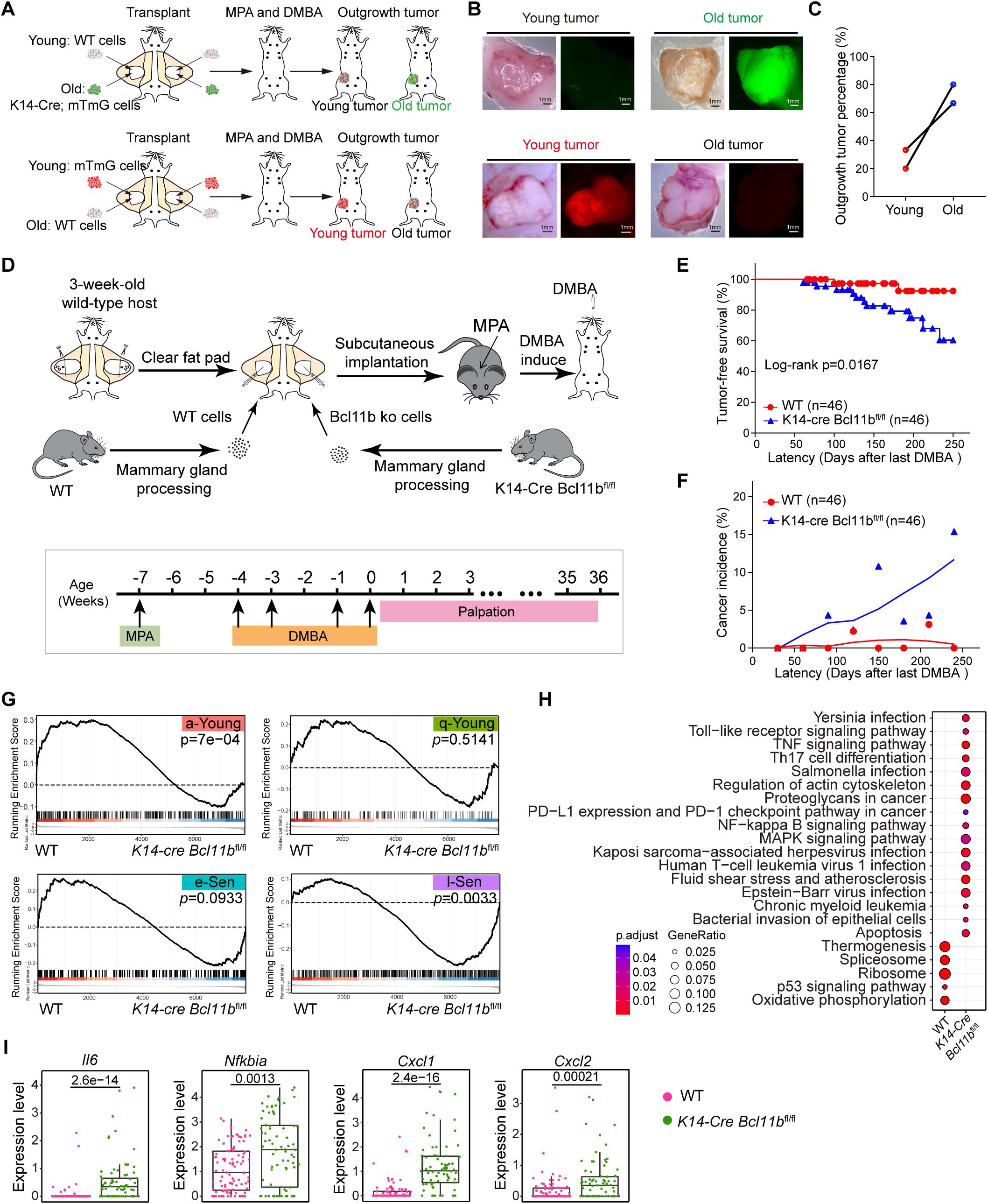
The l-Sen cells induced by knocking out *Bcl11b* expression were intrinsically susceptible to transformation into cancer cells. (A) Schematic diagram showing the DMBA induced cancer assay for young and old transplanted mammary cells. Mice transplanted with 2 month old young mammary cells (wt) and 24 month old mammary cells (with GFP reporter) mammary cells at equal MRUs were used for tumor formation assay. After cells grafted, mice were treated with MPA plus DMBA as indicated in the schematic diagram. Mice transplanted with 2 month old young mammary cells (with tdTomato reporter) with 12 month old mammary cells (wt) at equal MRUs were used for tumor formation assay. After cells grafted, mice were treated with MPA plus DMBA as indicated in the schematic diagram. (B) Representative images of tumors from young and old mammary gland cells. Scale bar, 1 mm. (C) Outgrowth tumor percentage of young and old mammary gland cells. (D) Schematic diagram showing the DMBA induced cancer assay for wild-type and *K14*-*Cre Bcl11b*^fl/fl^ transplanted mammary cells. Mice transplanted with WT or *K14*-*Cre Bcl11b*^fl/fl^ mammary cells were used for tumor formation assay. After cells grafted, mice were treated with MPA plus DMBA as indicated in the schematic diagram. (E) Tumor free survival curve of DMBA induced cancer assay for WT and *K14*-*Cre Bcl11b*^fl/fl^ transplanted mammary cells. Statistical analysis was determined by log-rank test. Latency was calculated from the day of last DMBA treatment. (F) Cancer incidence of WT and *K14*-*Cre Bcl11b*^fl/fl^ transplanted mice at each designated latency time (30 days for each period). (G) Gene Set Enrichment Analysis (GSEA) showing WT tumor cells were enriched for a-young signature genes and *K14*-*Cre Bcl11b*^fl/fl^ tumor cells were enriched for l-Sen signature genes. (H) KEGG pathway analysis of the differentially expressed genes in DMBA induced tumors in WT and *K14*-*Cre Bcl11b*^fl/fl^ CD49f^high^EpCAM^low^ cells. (I) SASP and NF-kB related genes were up-regulated in *K14*-*Cre Bcl11b*^fl/fl^ CD49f^high^EpCAM^low^ tumor cells.

### Multiple Aging- and Longevity-related Pathways Are Governed by *Bcl11b*

We then asked, how does *Bcl11b* play such a striking role in mammary cell senescence? To answer this question, we performed ChIP-seq analysis to profile the targets of *Bcl11b* in mammary cells. We detected 1197 Bcl11b binding sites in the genome, with 918 promoter binding sites and 279 enhancer binding sites (Figures 6A and 6B). The KEGG and GO pathway enrichment analysis revealed that many of the aging-associated pathways were the direct targets of Bcl11b, including the energy/nutrient sensing pathways ‘PI3K-Akt signaling’, ‘mTOR signaling’, ‘AMPK signaling’, which have been implicated in longevity regulation ^69^; the inflammation pathway ‘TNF signaling’; the fate determination pathway ‘Notch signaling’’, and ‘Wnt signaling’ and cancer-related pathways (Figures 6C and 6D). We also identified targets associated with pluripotency and the NF-kB pathway that were regulated by Bcl11b activity (Figure S9A). These pathways were also associated with both the q-Young-to-e-Sen and e-Sen-to-l-Sen cell transitions. These findings suggest that *Bcl11b* is a master regulator of aging progression by comprehensively repressing aging-associated pathways.

**Figure 6.**
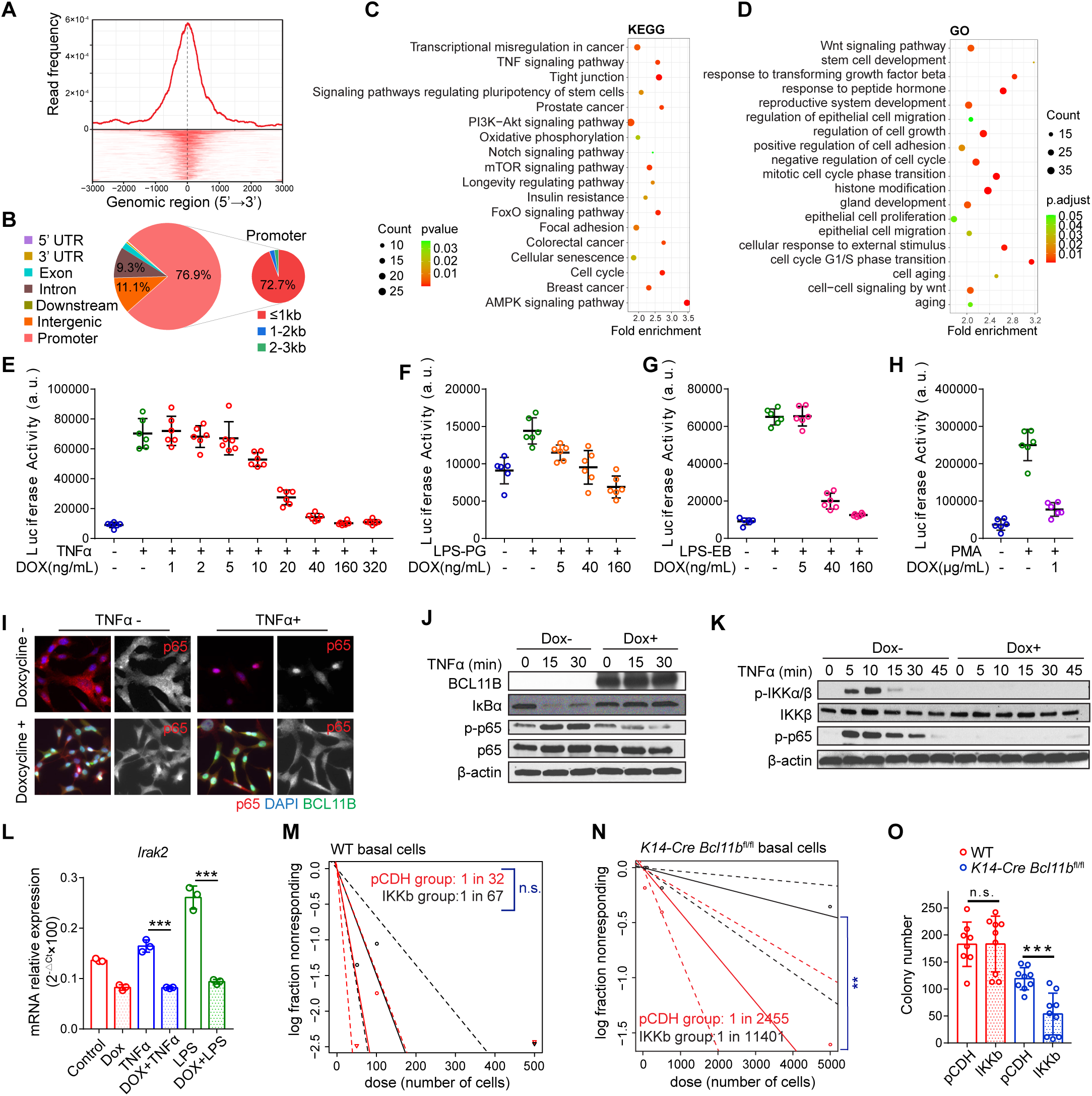
Multiple ageing associated pathways are governed by *Bcl11b*. (A) The distribution of global Bcl11b ChIP-seq peaks at transcription start site (TSS). (B) Pie chart showing Bcl11b ChIP-seq peak distribution in different genomic regions. (C-D) KEGG (C) and GO (D) pathway analysis of Bcl11b’s ChIP targets. (E-H) NF-kB luciferase reporter assay showing that *Bcl11b* expression regulates NF-kB activity. The pInducer-*Bcl11b* Comma Dβ cells with NF-kB luciferase reporter were treated with TNFα (E; 20ng/mL), LPS-PG (F; 1μg/mL), LPS-EB (G; 10^3^ EU/mL), PMA (H; 100ng/mL). In all the conditions, the induced expression of *Bcl11b* by doxycycline efficiently suppressed the activation of NF-kB pathway. (I) Immunofluorescence of p65 showing that induced expression of *Bcl11b* efficiently suppressed the nuclear import of p65 induced by TNFa (20ng/mL) in pInducer-*Bcl11b* Comma Dβ cells. Red: p65; Green: Bcl11b; Blue: DAPI. (J) Western blot analysis showing the induced Bcl11b expression by doxycycline (50ng/mL) inhibited IκBα degradation, p65 phosphorylation upon TNFα (20ng/mL) treatment in pInducer-*Bcl11b* Comma Dβ cells. (K) Western blot analysis showing the induced *Bcl11b* expression by doxycycline (50ng/mL) inhibited IKKa/b phosphorylation upon TNFα (20ng/mL) treatment in pInducer-*Bcl11b* Comma Dβ cells. (L) Real time PCR confirming *Irak2* mRNA expression is regulated by induced *Bcl11b* expression. N=3; data are presented as mean ± SD; two-tailed unpaired t-tests; ***P<0.001. (M) Extreme limiting dilution analysis (ELDA) plot showing the transplant of WT CD49f^high^EpCAM^low^ cells transduced with pCDH or pCDH-IKKb vectors. n.s., not significant. (N) ELDA plot showing the transplant of *K14*-*Cre Bcl11b*^fl/fl^ CD49f^high^EpCAM^low^ cells transduced with pCDH or pCDH-IKKb vectors. **P<0.01. (O) Colony formation assay of CD49f^high^EpCAM^low^ cells from WT and *K14*-*Cre Bcl11b*^fl/fl^ mice transduced with pCDH or pCDH-IKKb vectors. 3000 cells/well were seed and cultured for 1 week. n=9 per group; data are presented as mean ± SD; two-tailed unpaired t-tests; n.s., not significant, ***P<0.001.

As a stress sensing pathway, ‘NF-kB signaling’ has been recognized as an important contributor to the aging process; therefore, we wondered whether *Bcl11b* physiologically regulates NF-kB’s activity. A NF-kB luciferase activity assay suggested that induced expression of *Bcl11b* efficiently repressed NF-kB activity that had been triggered by TNFα, lipopolysaccharide (LPS) or phorbol myristate acetate (PMA) (Figures 6E-6H). *Bcl11b* regulates NF-kB expression by upstream of the signaling pathway, as indicated by the nuclear transport of RelA, the degradation of IkBa, and the activation of IKKa/b all blocked by *Bcl11b* expression (Figures 6I-6K and S9B). When we analyzed our ChIP-seq data, we found that Bcl11b directly bound to the promoter regions of *Irak2* (an essential NF-kB signaling mediator), *Nfkbia* and *Nfkb2*, suggesting direct transcriptional regulation (Figure S9A). Indeed, when we induced the expression of *Bcl11b*, the mRNA expression of *Irak2* was significantly suppressed (Figures 6L and S9C). These data collectively demonstrate that *Bcl11b* directly regulates the mammary cell stress response program as one of the mechanisms to slow aging progression.

### NF-kB Promotes Stem Cell Exhaustion in the Absence of *Bcl11b*

NF-kB signaling is a well-characterized stress-sensing pathway ^70^ and plays pleiotropic roles in a variety of biological processes ^71^. This pathway is aberrantly activated during aging through an unclear mechanism. We asked, under what conditions does NF-kB activation convert a cell from a young state to a senescent state? To answer this question, we first tested stem cell activity after NF-kB activation in WT cells and *Bcl11b*-KO cells and found that the enforced expression of IKKb triggered NF-kB activation (Figures S10A-10C) but did not significantly reduce the mammary reconstitution ability in the WT cells (1 in 32 of the control cells vs. 1 in 67 of the IKKb-overexpressing cells, p>0.05) (Figure 6M, S10D, S10F, S10I and S10J). However, when the activation of NF-kB was performed in cells with a *Bcl11b*-KO background, the mammary reconstitution ability was significantly reduced (1 in 2455 *Bcl11b*-KO cells vs. 1 in 11401 *Bcl11b*-KO+IKKb-positive cells, p<0.01) (Figures 6N, S10E, S10G, S10I, S10K and S10L). The differential NF-kB activation effects in cells with different backgrounds imply that NF-kB activity alone is not sufficient to drive stem cell exhaustion, and the reduced regeneration ability, which is a frequent hallmark of tissue aging, is initiated through a specific epigenetic modification program involving multiple signaling pathways that depend on *Bcl11b* expression. In accordance with this idea, we observed differential colony formation rates of IKKb-expressing WT and *Bcl11b*-KO cells (Figures 6O and S10H), indicating that *Bcl11b* may protect mammary cells from aging in response to NF-kB activation.

### Manipulation of l-Sen Cells Regulates Aging-related Cancer Transformation

We then explored the possibility of reversing the progressive aging process as a strategy to reduce cancer vulnerability. As the *Bcl11b* activity declined with age, we screened a small customized drug pool to identify a chemical drug that would enable us to reduce the number of senescent cells and increase the number of young cells. We tested PI3K-Akt-mTOR inhibitors, NF-kB pathway inhibitors, Jak-Stat inhibitors, wnt inhibitors, notch inhibitors and hedgehog inhibitors to evaluate their roles in enhancing *Bcl11b* expression, which is an indication of a young cell state. We found that TPCA-1 ^72,73^, a dual NF-kB and Jak-Stat inhibitor, exerted the most striking effect in restoring *Bcl11b* expression, while other NF-kB or Jak-Stat single pathway inhibitors played minimal roles in *Bcl11b* expression, indicating that aging is a progressive and coordinated process that might require the input of multiple pathways (Figures 7A and S11A). Consistent with the role played by TPCA-1 role in promoting *Bcl11b* expression in vitro (Figures 7B and S11B), when we administered TPCA-1 to 12-month-old mice continuously for 1 month, mammary CD49f^high^EpCAM^low^ cells were clearly younger, showing a remarkable increase in a-Young cells accompanied by a dramatic reduction in the number of e-Sen and l-Sen cells (Figures 7C, 7D and S11C-S11I). The *Bcl11b* activity score and metabolic pathways were clearly restored after TPCA-1 treatment, while aging-related pathways, including the senescence, NF-kB, Jak-Stat, MAPK and PI3K-Akt pathways, were all significantly suppressed (Figures 7E-7H). This molecular profile is very similar to that of the 2-month-old mammary glands, suggesting that TPCA-1 can change the senescence profile of mammary gland cells under physiological conditions. To test whether TPCA-1’s rejuvenation effect is dependent on Bcl11b expression, we performed TPCA-1 treatment on 3-month-old Bcl11b KO mice continuously for 1 month, we found that the ageing phenotype of Bcl11b KO mammary cells on the pseudotime can be efficiently rescued by TPCA-1 treatment (Figure S12). The l-Sen cells of TPCA-1 treated mice were significantly reduced compared with Bcl11b KO cells, and the SASP pathway, NFkb signaling pathway, JAK-STAT signaling pathway were all efficiently suppressed. This suggests that when Bcl11b’s downstream target signaling pathways were inhibited, it can play a similar role as the Bcl11b expression and block the accelerated ageing progression.

**Figure 7.**
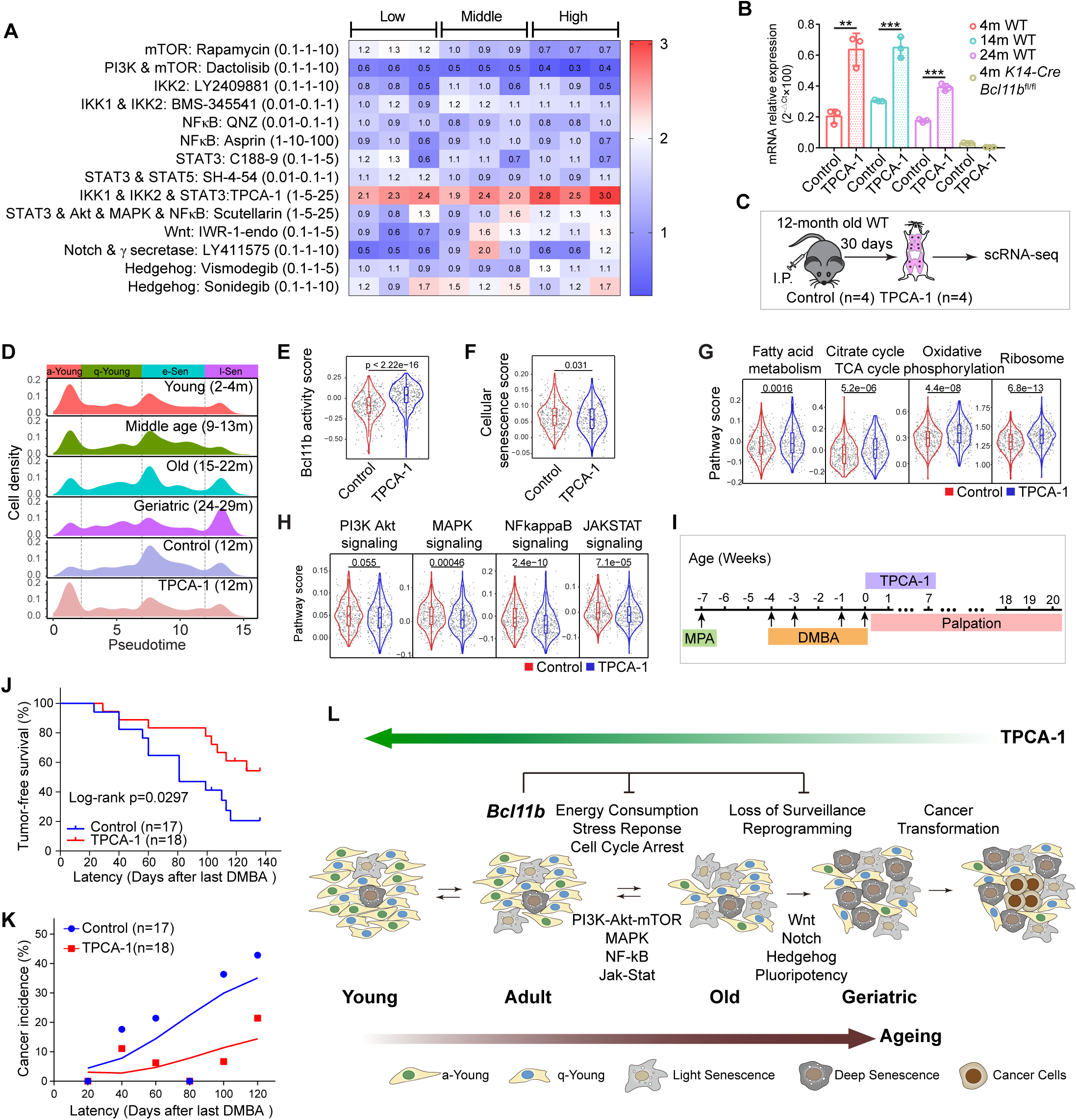
TPCA-1 reshapes ageing programs at the transcriptome level. (A) Evaluation of drugs that can efficiently upregulate *Bcl11b* expression. Real time PCR quantification of *Bcl11b* expression levels treated with low-middle-high dose of Rapamycin, Dactolisib, LY2409881, BMS-345541, QNZ, Asprin, C188-9, SH-4-54, TPCA-1, Scutellarin, IWR-1-endo, LY411575, Vismodegib and Sonidegib with β-actin as an internal control (n=3) in Comma Dβ cells. Fold changes compared to the control group were shown in the data matrix. Data were compiled and presented as heatmap. (B) Real time PCR confirming *Bcl11b* mRNA expression were upregulated by TPCA-1 (25μM) treatment in primary mammary CD49f^high^EpCAM^low^ cells isolated from 4m WT, 14m WT, 24m WT, 4m *K14*-*Cre Bcl11b*^fl/fl^ mice. Basically, CD49f^high^EpCAM^low^ cells from 4-month-old, 14-month-old, 24-month-old WT mice and 4-month-old *K14*-*Cre Bcl11b*^fl/fl^ mice were sorted into the 96-well Ultra-Low attachment culture plate and cultured with 25μM TPCA-1 or DMSO for 24h. Live cells were subsequently sorted out for RT-PCR assay. n=3; bar and whiskers denote mean ± SD; two-tailed unpaired t-tests; **P<0.01, ***P<0.001. (C) Schematic diagram showing the strategy of TPCA-1 (10mg/kg) treatment in vivo on 12-month-old mice. (D) Cell density distribution of mammary CD49f^high^EpCAM^low^ cells from young, middle age, old, geriatric mice, along with control (12-month-old, n=4) and TPCA-1treated (12-month-old, n=4) mice over pseudotime. (E-F) Violin plots showing the increased *Bcl11b* activity score (E) and decreased cellular senescence score (F) in TPCA-1 treated group. (G-H) KEGG analysis of the up-regulated (G) and down-regulated (H) pathways in CD49f^high^EpCAM^low^ cells of TPCA-1 treated mammary gland. (I) Schematic diagram showing the DMBA induced cancer assay for mice treated with DMSO (4%) and TPCA-1 (10mg/kg). Mice were treated with MPA plus DMBA followed by TPCA-1 as indicated in the schematic diagram. (J) Tumor free survival curve of DMBA induced cancer assay for mice treated with DMSO (4%) and TPCA-1 (10mg/kg) as indicated. Statistical test was performed by log-rank test. Latency was counted from the date of the last DMBA treatment. (K) Cancer incidence of DMSO (4%) and TPCA-1 treated mice at each designated latency time (20 days for each period). Latency was counted from the date of the last DMBA treatment. (L) Schematic diagram showing the hypothesis of how the progressive senescence programs result in cancer vulnerability.

To determine whether TPCA-1 induced younger mammary gland can efficiently reduce cancer incidence, we performed 7 weeks of intraperitoneal administration of TPCA-1 after medroxyprogesterone acetate (MPA)- and DMBA-treatment, and tracked the rate of cancer incidence (Figure 7I). We found that TPCA-1 treatment successfully and efficiently decreased the tumor burden and significantly increased tumor-free mouse survival (Figures 7J and 7K). These data demonstrate that mammary cancer formation can be controlled with a strategy that reshape the aging tissue transcriptome to that of tissue in the juvenile state to reduce cancer susceptibility.

## DISCUSSION

The global aging trend has become increasingly common worldwide, with an anticipated increase in cancer burden. The biological relationship between aging and cancer has been a critical issue to clarify to guide prophylactic measures. Our chronological single-cell transcriptome analysis of the mammary gland enabled us to reconstruct a molecular portrait of the physiological aging process, which revealed heterogeneous senescent cell states and progressive aging processes intrinsic to epithelial cells. This molecular map bridges cellular senescence and cancer initiation and answers the long-standing question, why do aging cells with degenerative activities paradoxically foster cancer formation? Our study implies that the senescent state is an entropic arrested cell state with aberrantly activated stem cell and cancer programs that are modulated by the *Bcl11b*-associated signaling network. The paradox can be explained by a progressive ageing model that senescence and cancer are successive biological steps in the same linear developmental trajectory, not bifurcating biological processes (Figure 7L). This understanding can help us design novel strategies to block aging and cancer at various intersections.

Our study led us to rethink the prevailing mutation accumulation model for explaining the aging– cancer relationship. In the last century, Carl Nordling proposed the theoretical framework that carcinogenesis is driven by mutation of the genome ^74^. This seminal concept was first developed into a multistage model for malignancy transformation ^75^. However, this theory does not explain why a substantial portion of mutations occur early in life, while cancers arise exponentially later in life ^76,77^. Neither does it explain the disproportion between cancer frequency and animal body size, as well as the scaling of cancer incidence to animal lifespan ^78^. These facts implicate that there might be more factors beyond genetic mutations involved in determination of cancer initiation. In this study, we clearly observed transcriptome alterations during aging, and these were biologically correlated with cancer initiation, and we identified a master fate determinant, *Bcl11b*, which is involved in epigenetic regulation ^79^ in aging and cancer. In addition, we demonstrated that after mutation, modulation of epigenetic reprogramming with chemical inhibitors targeting NF-kB and Jak-stat efficiently reduced cancer formation. This outcome suggests that although mutation is essential and may be indispensable for triggering cancer formation, epigenetic programs might be equally crucial to cancer development and may be more manageable. Therefore, modulating epigenetic modifications may be a better and more feasible way than genetic intervention to control cancer incidence at the population level. We envision that a comprehensive molecular understanding of epigenetic regulation in aging may help us find diverse and promising targets to eventually reduce cancer occurrence.

Meanwhile, our study has limitations at that we investigated a specific cell population in the mammary gland in the longitudinal ageing process and we utilized DMBA treatment-based cancer initiation model to establish the molecular connections between ageing and cancer. It will be interesting and promising to test in the future whether the ageing dynamics we discovered hold true for other mammary populations or even other organs in a variety of cancer models.

---

**Figure S1.**
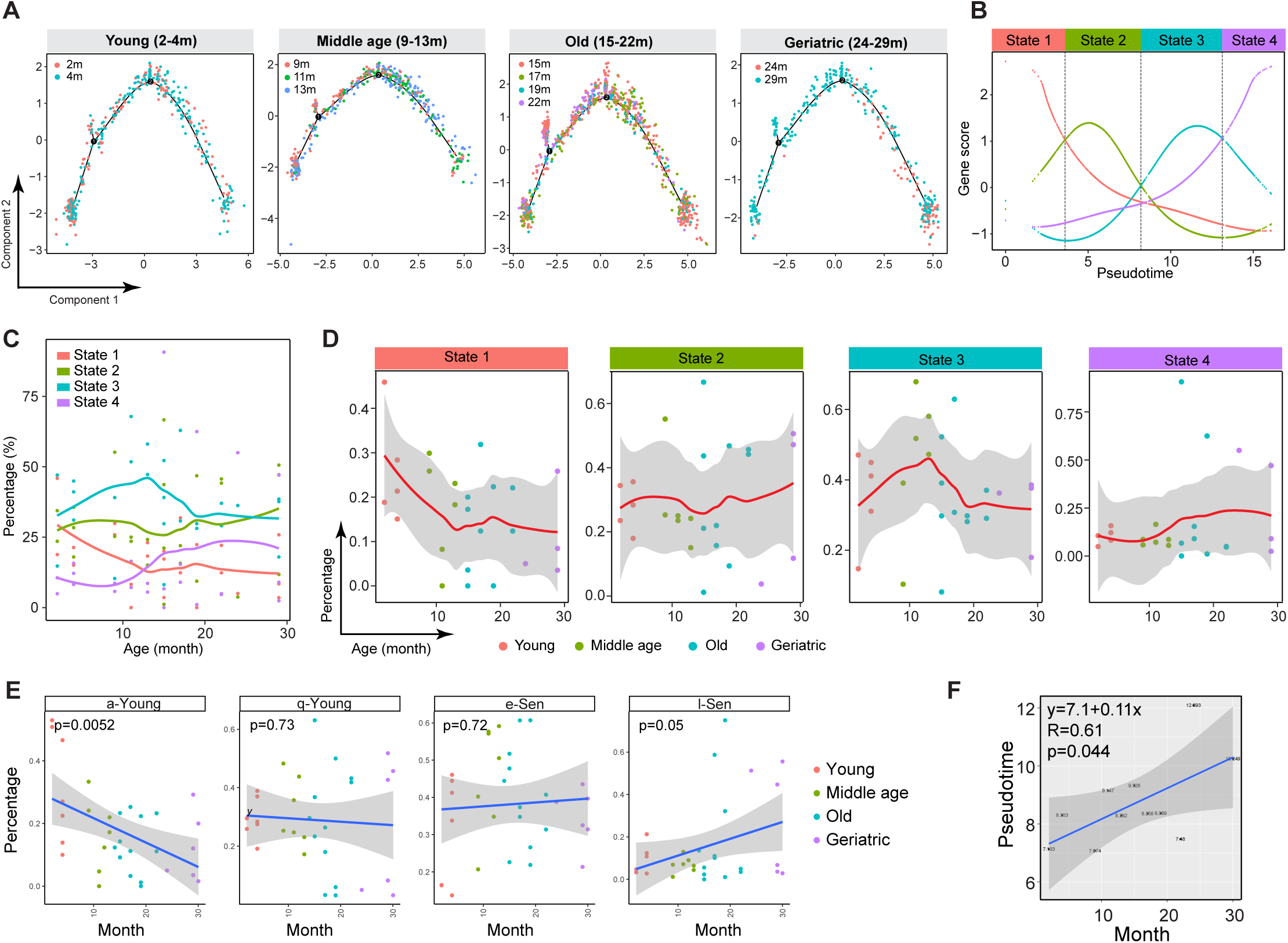
Distribution of mammary cells over pseudotime trajectory and age progression. (A) Pseudotemporal ordering of mammary cells from various age groups showing that each age group contains all the cell states. Cells are labeled by colors according to their biological ages in each age group. (B) Expression dynamics of the signature gene clusters of each cell state over the pseudotime trajectory. The boundary of each cell state was determined by the intersections of gene scores from neighboring cell states. (C-D) Dynamics of the proportion of each cell state over the real chronological age. (E) The statistics of the dynamics of various mammary cell states. (F) The correlation between mouse age and pseudotime.

**Figure S2.**
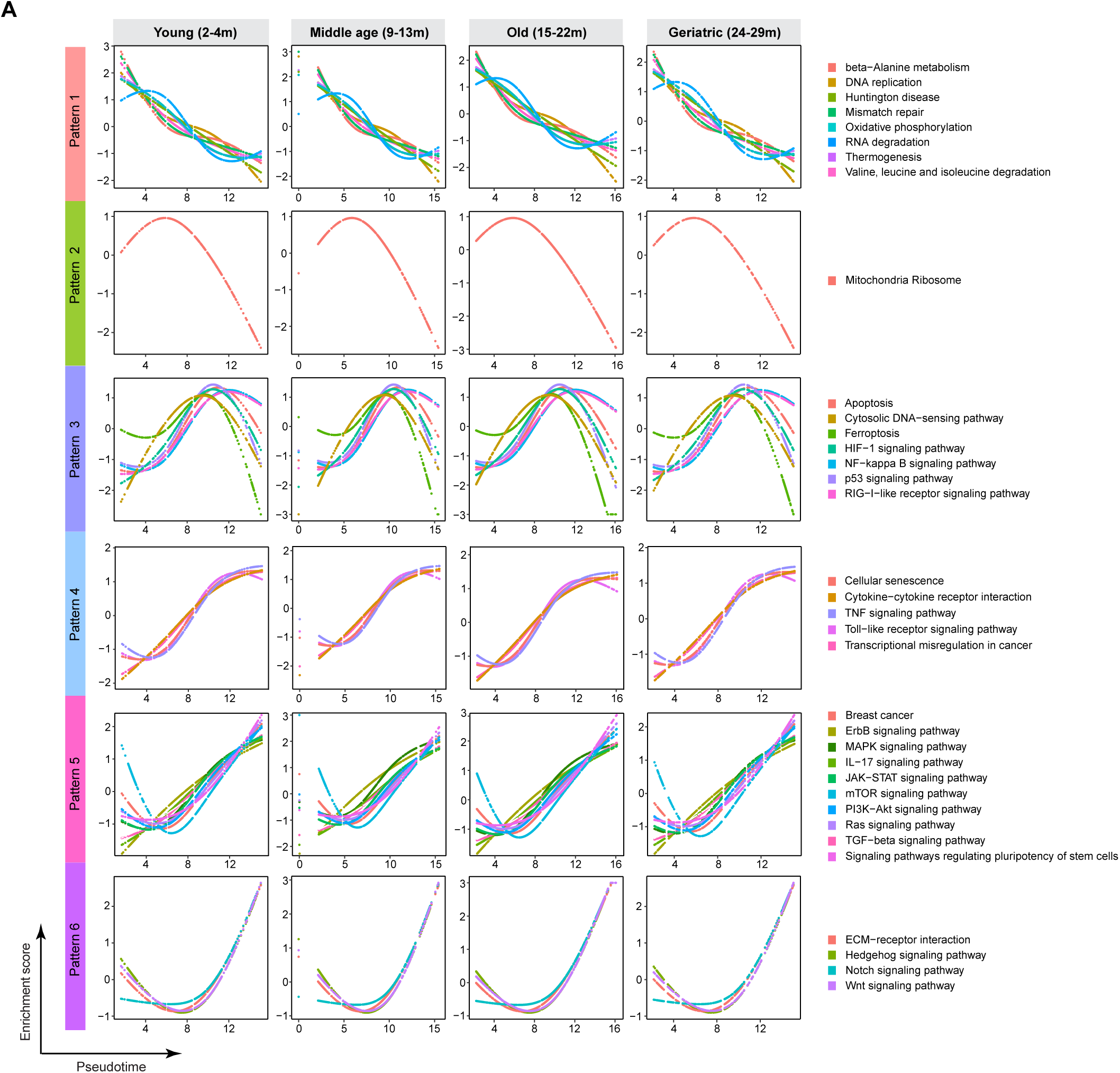
Pathway dynamics of mammary cells from each age group over the pseudotime. (A) Enrichment score of pathways from 6 different patterns for young, middle age, old and geriatric mammary cells. Note: the dynamic pattern of each signaling pathway for mammary cells from different age group remains largely the same. This indicates that the dynamic pattern of signaling pathways are determined by cell state instead of the real chronological age.

**Figure S3.**
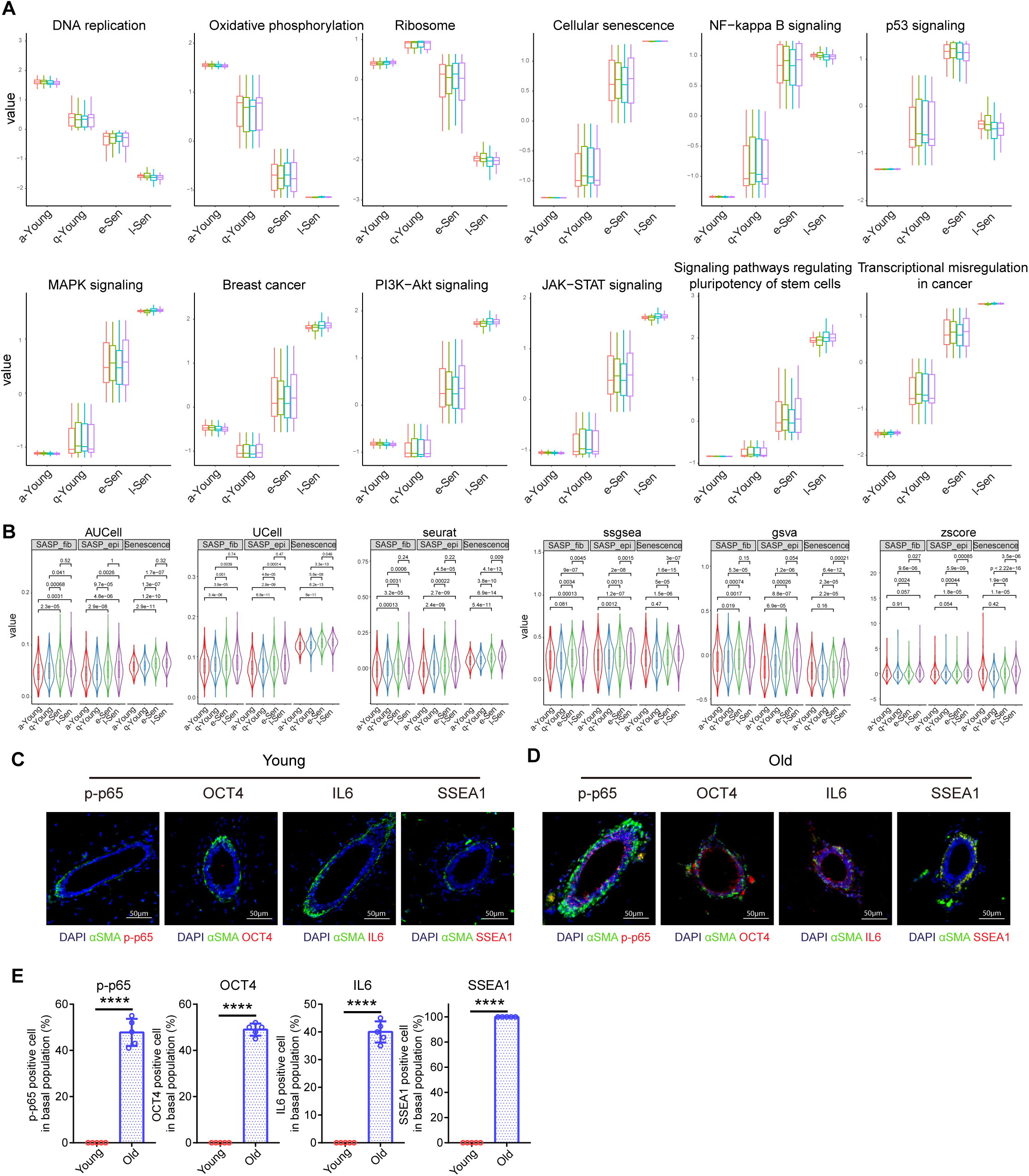
The aging related pathways in various mammary cell states and mouse ages. (A) Signaling pathway analysis within cellular states in various mouse ages. (B) The aging related pathways in various mammary cell states. (C) Representative immunofluorescence staining of p-p65, Oct4, Il6 and Ssea1 (scale bar, 50μm) in mammary cells from young mice. (D) Representative immunofluorescence staining of p-p65, Oct4, Il6 and Ssea1 (scale bar, 50μm) in mammary cells from old mice. (E) Quantification of p-p65, Oct4, Il6 and Ssea1 in young and old mammary glands. Statistical analysis was performed using two-tailed unpaired t-tests; data are presented as mean ± SD; ****P<0.0001.

**Figure S4.**
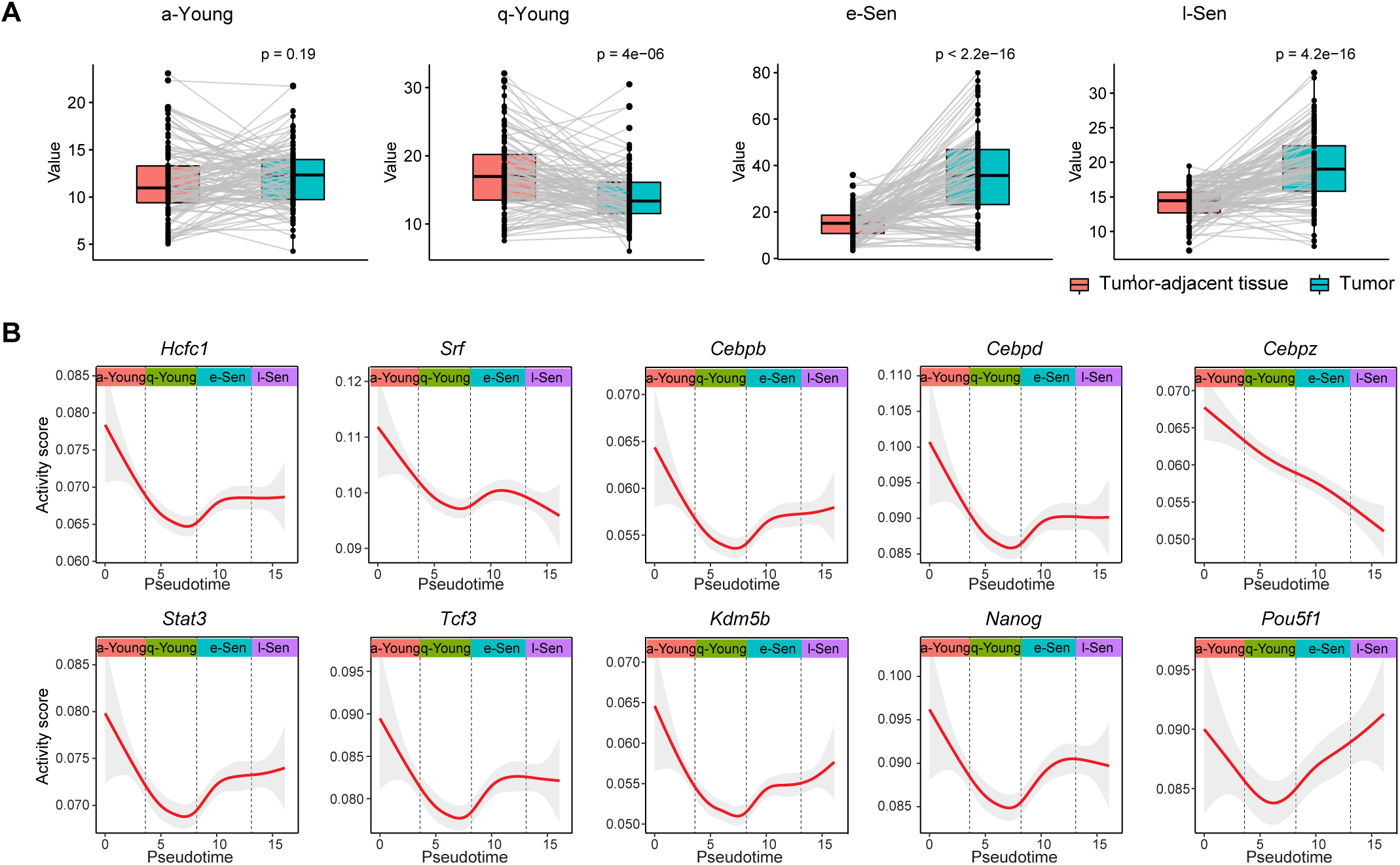
Activity score of transcription factors enriched in a/q-Young and l-Sen states. (A) Cell state signature in tumor-adjacent tissue and tumor tissues. (B) Activity score of transcription factors enriched in a/q-Young and l-Sen states presented by target gene score over the pseudotime. The aging related pathways in various mammary cell states.

**Figure S5.**
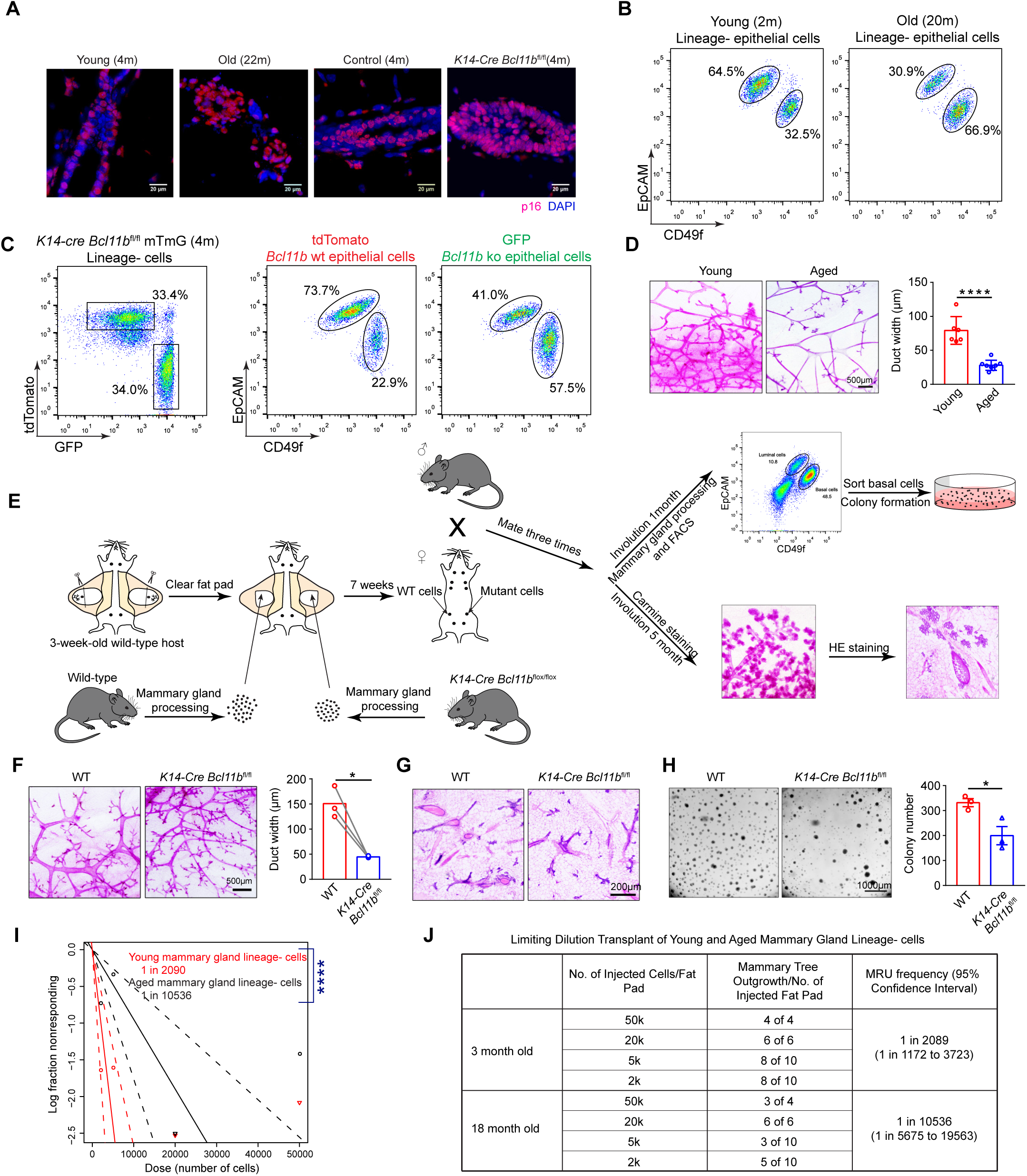
Accelerated stem cell exhaustion during ageing and after *Bcl11b* knockout. (A) Representative immunofluorescence staining of p16^Ink4a^ (scale bar, 20μm) in mammary epithelial cells from young (4 month), old (22 month), control *K14*-*Cre Bcl11b*^wt/wt^ (4 month) and *K14*-*Cre Bcl11b*^fl/fl^ (4 month) mice. (B) Representative FACS plots of basal cells and luminal cells in young (2m) and old (20m) mammary gland lineage-population. (C) FASC analysis of mammary epithelia in *K14*-*Cre Bcl11b*^fl/fl^ *mTmG* mice showing the relative proportion of basal cells and luminal cells. (D) Representative wholemount staining and duct width quantifications of young (2m-5m) and aged (12m-29m) mammary glands. Scale bar, 500μm; statistical analysis was performed using two-tailed unpaired t-tests; data are presented as mean ± SD; ****P<0.0001. (E) Schematic diagram for the workflow of the ageing phenotype quantification for WT and *K14*-*Cre Bcl11b*^fl/fl^ mammary cells after multiple rounds of reproduction cycles. Basically, WT and *K14*-*Cre Bcl11b*^fl/fl^ mammary cells were transplanted onto cleared fat pads. Recipient mice were mated with male mice for 3 rounds. Whole mount and colony formation assays were performed after the last round of pregnancy. (F-H) Representative pictures of mammary wholemounts (F; Scale bar, 500μm), HE staining (G; Scale bar, 200μm) and colony formation (H; Scale bar, 1000μm) of WT and *K14*-*Cre Bcl11b*^fl/fl^ transplanted mammary glands after multiple rounds of pregnancy. Statistical analysis was performed using two-tailed unpaired t-tests; data are presented as mean ± SD; *P<0.05. (I) ELDA plot of limiting dilution transplant of young (3m) and old (18m) mammary gland lineage-cells. P<0.0001. (J) Table for limiting dilution transplant of young (3month) and old (18 month) mammary gland lineage-cells.

**Figure S6.**
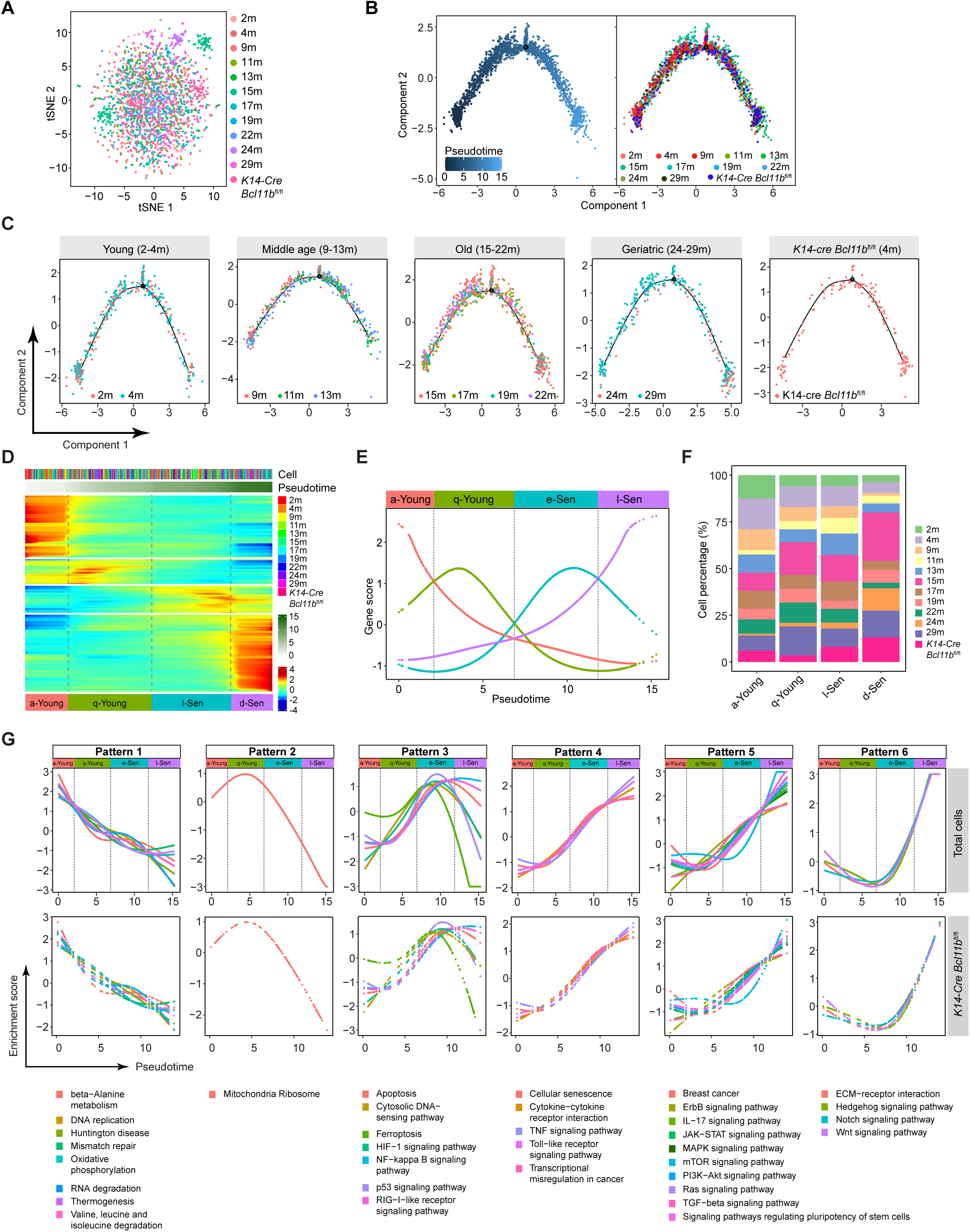
Single cell ageing trajectory analysis of WT (2-29 months) with *Bcl11b* KO CD49f^high^EpCAM^low^ cells (4months). (A-B) t-SNE plot showing the clustering analysis (A) and the pseudotime ordering analysis (B) of scRNA-seq data from young, middle age, old, geriatric mammary cells along with *K14*-*Cre Bcl11b*^fl/fl^ (4month) CD49f^high^EpCAM^low^ cells. Cell origins are labeled by colors. (C) Pseudotemporal ordering of mammary cells from various age groups showing that each age group contains all the cell states. Cells of different ages are labeled by colors in each age group. (D) Heatmap visualization of the dynamic gene expression changes over the pseudotime. (E) Score of differentially expressed genes in each re-analyzed stage along the pseudotime. Definition of a state is based on the intersection point of the neighboring cell states. (F) Relative proportion of mammary cells from various ages of WT mice and *Bcl11b* KO (4month) in each re-analyzed cell state. (G) Activity score of each signaling pathway over the pseudotime of WT mammary cells and *K14-Cre Bcl11b*^fl/fl^ cells. Note: the dynamic pattern of each signaling pathway for mammary cells from different age group and *K14*-*Cre Bcl11b*^fl/fl^ group remains largely the same.

**Figure S7.**
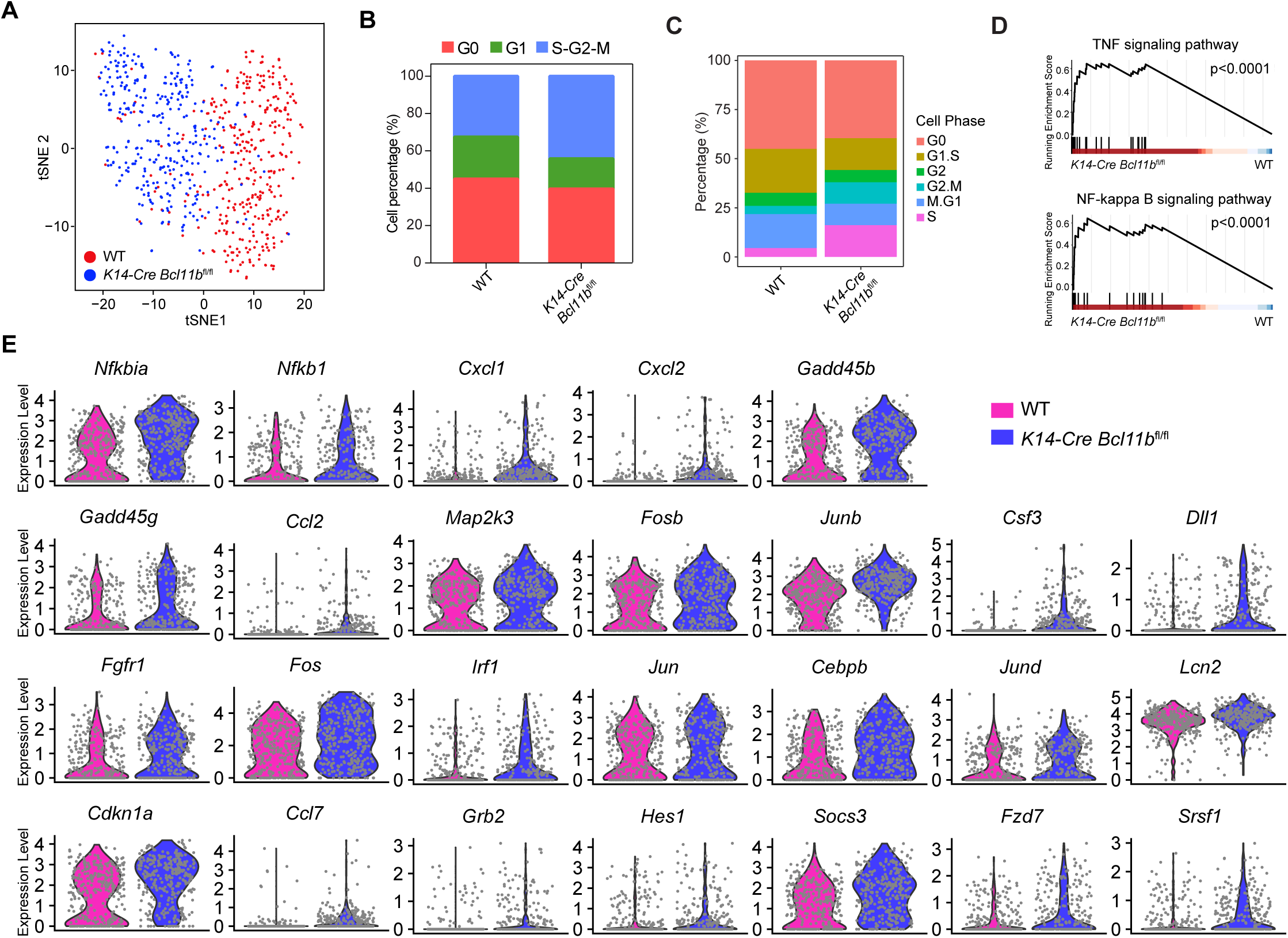
Differentially expressed genes in WT and *K14*-*Cre Bcl11b*^fl/fl^ CD49f^high^EpCAM^low^ mammary cells. (A) t-SNE plot showing the scRNA-seq data from WT (red; 4m) and *Kr14*-*Cre Bcl11b*^fl/fl^ (blue; 4m) CD49f^high^EpCAM^low^ mammary cells. (B) Cell cycle fraction analysis of G0, G1, S-G2-M in WT and *K14*-*Cre Bcl11b*^fl/fl^ groups. (C) Relative cell cycle proportion in WT and *K14*-*Cre Bcl11b*^fl/fl^ groups. (D) Gene set enrichment analysis (GSEA) enrichment analysis of scRNA-seq data from WT and *K14*-*Cre Bcl11b*^fl/fl^ CD49f^high^EpCAM^low^ cells showing the enrichment of TNF and NF-kB signaling pathways in *Bcl11b* KO mammary cells. (E) Differentially expressed genes in WT and *K14*-*Cre Bcl11b*^fl/fl^ CD49f^high^EpCAM^low^ cells.

**Figure S8.**
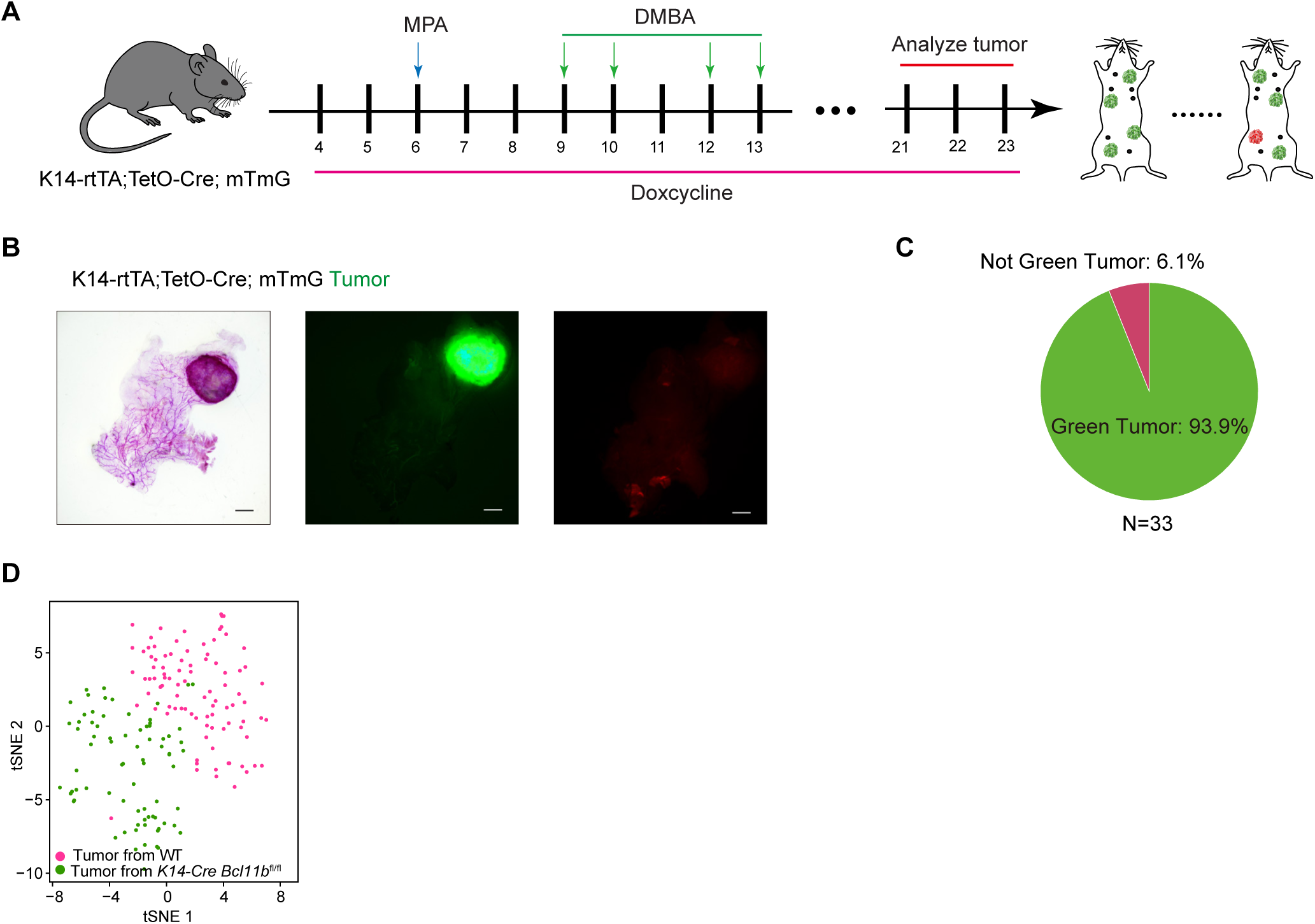
DMBA induced mouse breast tumors mainly originate from basal cells. (A) Basal cells of Krt14rtTA-TetOcre-mTmG mice were labeled with doxycycline induction, and mice were treated with DMBA. (B) Representative tumor images of Krt14rtTA-TetOcre-mTmG mice induced with doxycycline. (C) Green tumor percentage in the outgrowth tumors (N=33). (D) The t-SNE plot showing the clustering of CD49f^high^EpCAM^low^ cells from DMBA induced WT or *K14*-*Cre Bcl11b*^fl/fl^ tumor.

**Figure S9.**
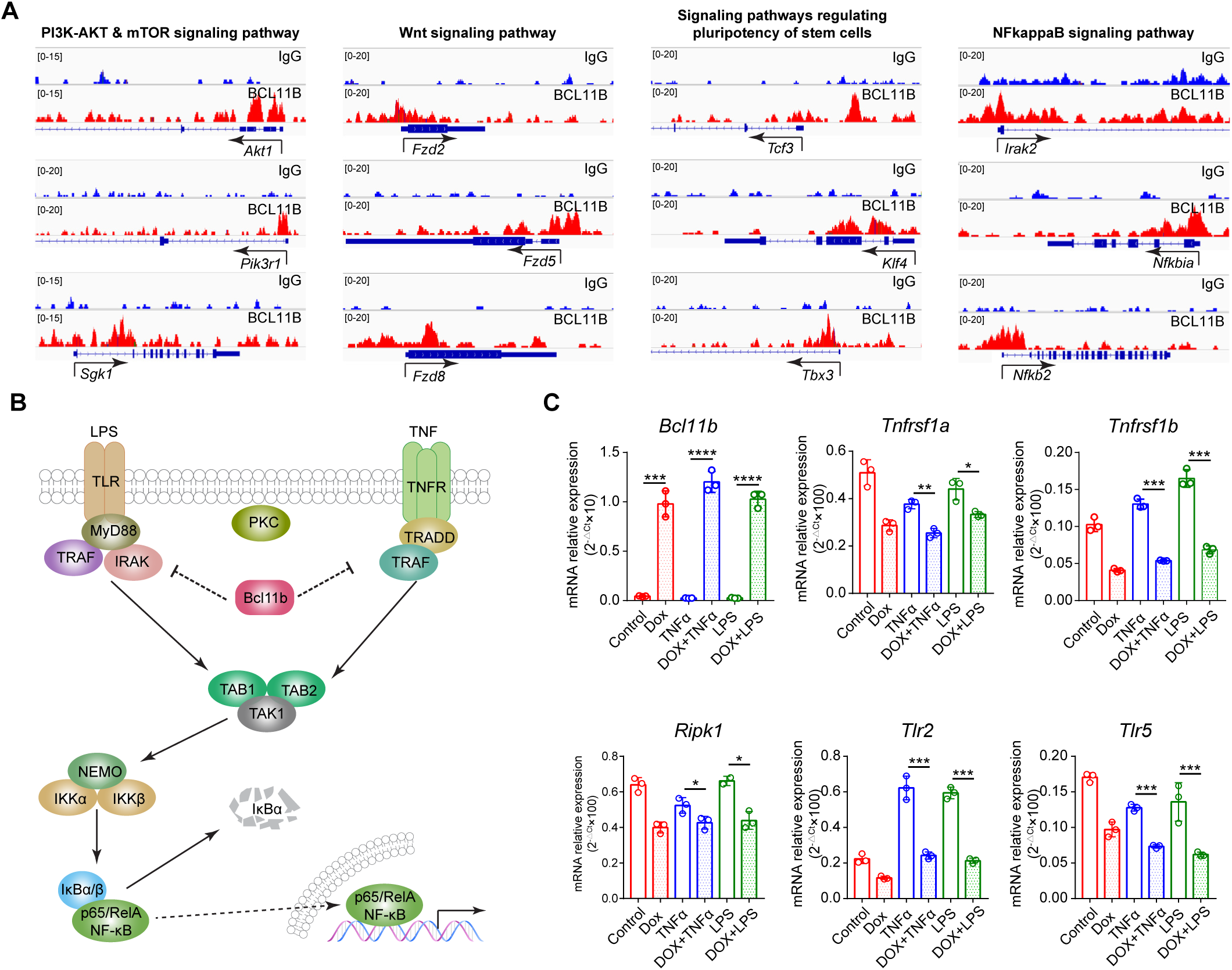
***Bcl11b* regulates genes involved in NF-kB signaling.** (A) ChIP-seq tracks showing Bcl11b ChIP binding targets in “PI3K-AKT & mTOR signaling pathway”, “Wnt signaling pathway”, “Signaling pathways regulating pluripotency of stem cells” and “NFkappaB signaling pathway” The exon/intron diagram is shown at the bottom and arrowheads represent the direction of transcription. (B) Schematic diagram showing the putative regulation of NF-κB pathway by *Bcl11b*. (C) Real time PCR analysis of NF-κB pathway related gene expression after induction of *Bcl11b* expression. N=3; bar and whiskers denote mean ± SD; two-tailed unpaired t-tests; *P<0.05, **P<0.01, ***P<0.001.

**Figure S10.**
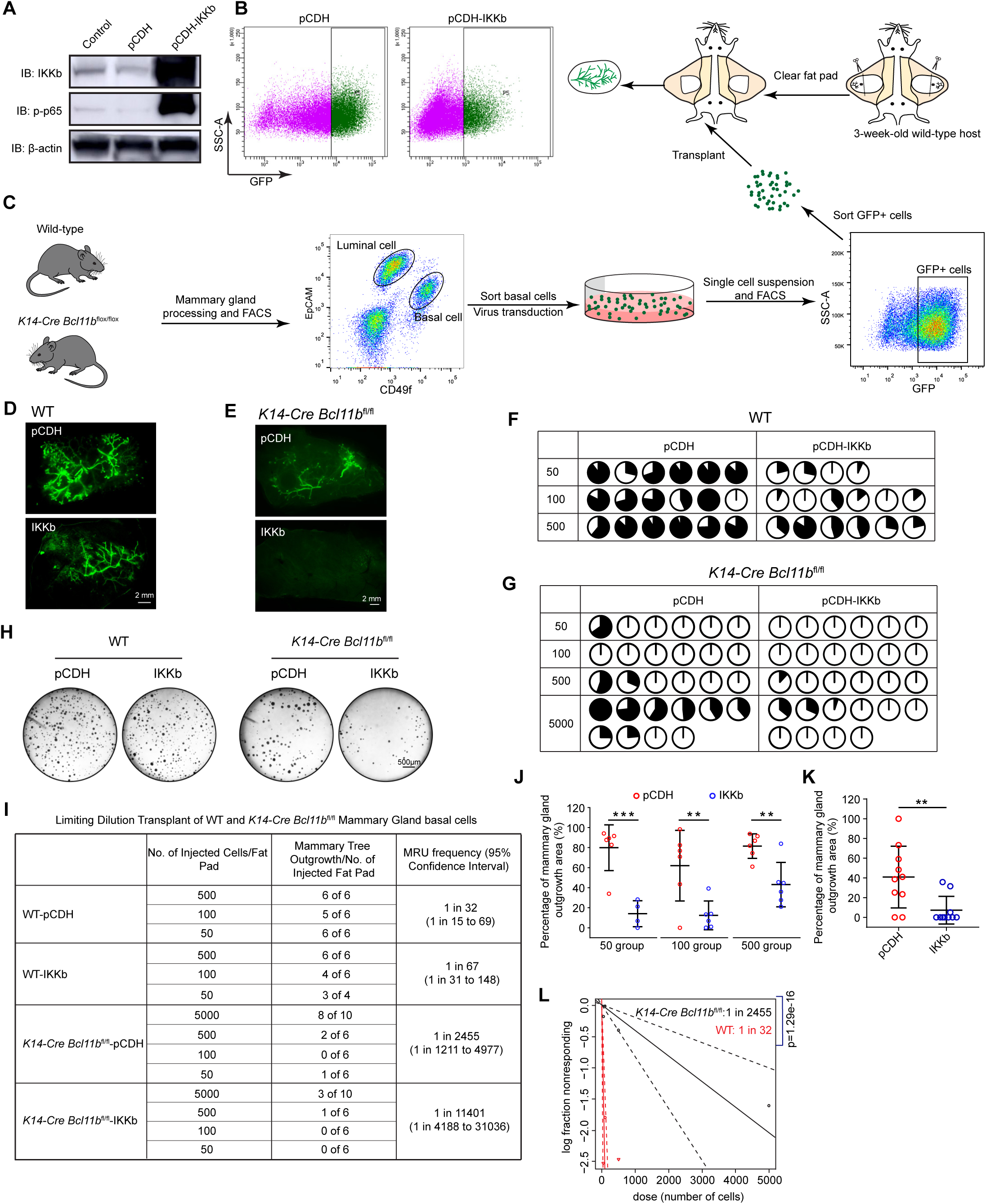
NF-kB activation by IKKb overexpression drives stem cell exhaustion in the absence of *Bcl11b*. (A) Western blot analysis of IKKb and p-p65 expression in 293T cells upon enforced IKKb expression. (B) FACS plot showing primary CD49f^high^EpCAM^low^ cells transduced with pCDH or pCDH-IKKb vectors. (C) Experimental schematic diagram for limiting dilution transplants of WT and *K14*-*Cre Bcl11b*^fl/fl^ CD49f^high^EpCAM^low^ cells transduced with pCDH or pCDH-IKKb vectors. (D) Wholemount images of representative outgrowths of WT CD49f^high^EpCAM^low^ cells transduced with pCDH or pCDH-IKKb vectors. Scale bar, 2 mm. (E) Wholemount images of representative outgrowths of *K14*-*Cre Bcl11b*^fl/fl^ CD49f^high^EpCAM^low^ cells transduced with pCDH or pCDH-IKKb vectors. Scale bar, 2 mm. (F) Pie chart of transplant of WT CD49f^high^EpCAM^low^ cells transduced with pCDH or pCDH-IKKb vectors. n.s., not significant. (G) Pie chart of transplant of *K14*-*Cre Bcl11b*^fl/fl^ CD49f^high^EpCAM^low^ cells transduced with pCDH or pCDH-IKKb vectors. **P<0.01. (H) Representative colony formation assay images of CD49f^high^EpCAM^low^ cells from WT and *K14*-*Cre Bcl11b*^fl/fl^ mice transduced with pCDH or pCDH-IKKb vectors. 3000 cells/well were seed and cultured for 1 week. Scale bar, 500μm. (I) Table for limiting dilution transplant of WT and *K14*-*Cre Bcl11b*^fl/fl^ mammary gland CD49f^high^EpCAM^low^ cells transduced with pCDH or pCDH-IKKb vectors. (J-K) Quantification of outgrowth area as proportion of total mammary gland area in WT (J) and *K14*-*Cre Bcl11b*^fl/fl^ (K) groups. Statistical analysis was performed using two-tailed unpaired t-tests; bar and whiskers denote mean ± SD; **P<0.01, ***P<0.001. (L) ELDA plot showing the comparison of WT and *K14*-*Cre Bcl11b*^fl/fl^ CD49f^high^EpCAM^low^ cells transduced with pCDH vectors.

**Figure S11.**
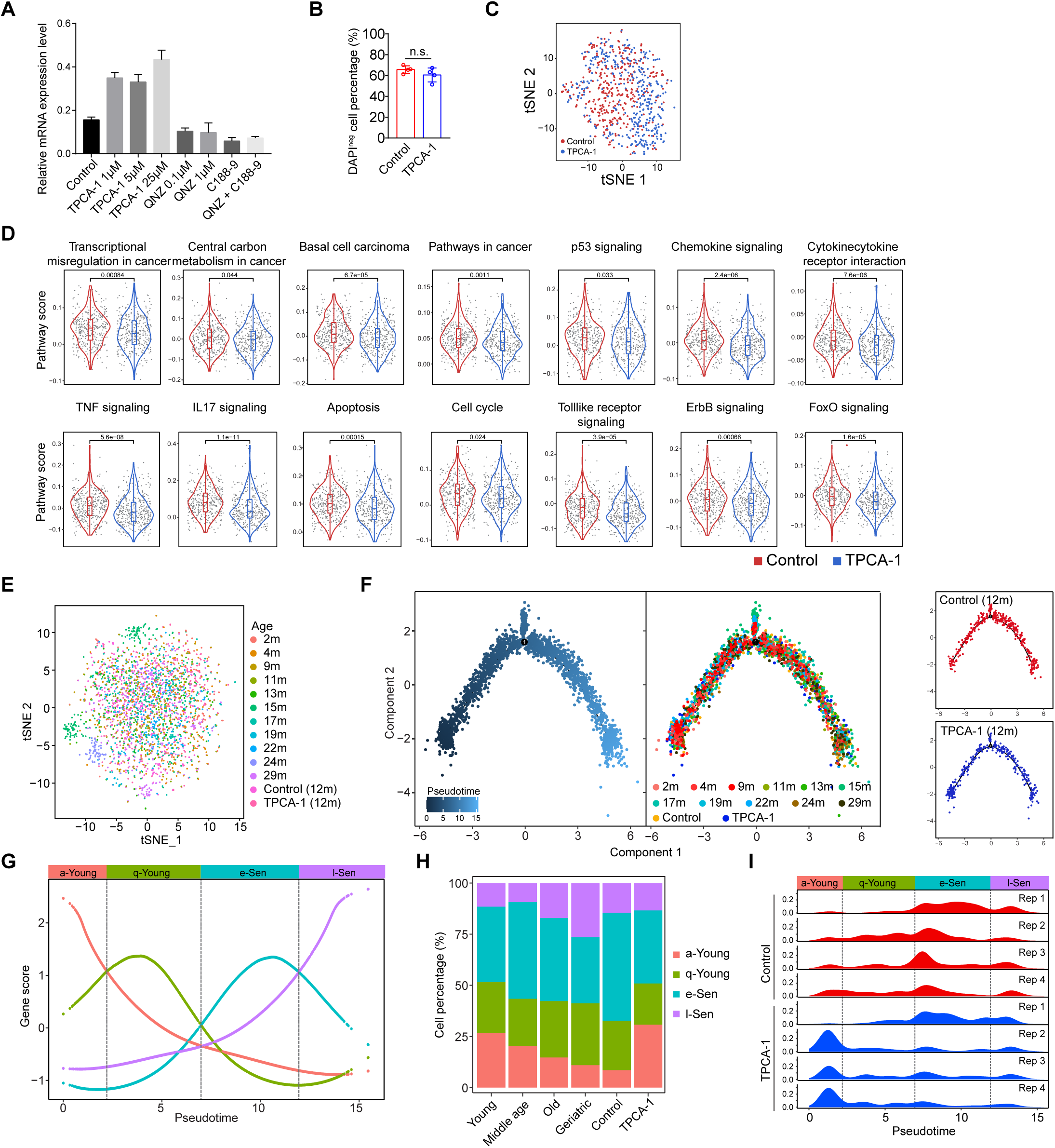
TPCA-1 reshapes ageing programs at the transcriptome level. (A) Real time PCR quantification of *Bcl11b* expression levels treated with QNZ, C188-9 and TPCA-1 with β-actin as an internal control (n=3) in Comma Dβ cells. (B) Percentage of DAPI negative cells in primary basal cells treated with TPCA-1. n.s., not significant. (C) tSNE plot showing the clustering of CD49f^high^EpCAM^low^ cells from 12m-old month WT mice with or without TPCA-1 treatment. (D) Down-regulated signaling pathways upon TPCA-1 treatment. (E) tSNE plot showing the clustering of CD49f^high^EpCAM^low^ cells from different age WT mice and 12m-old month WT mice with or without TPCA-1 treatment. (F) Pseudotime-ordering analysis of single cell transcriptomes from WT mice of various ages and 12m-old month WT mice with or without TPCA-1 treatment. (G) Cell state signature gene scores over the pseudotime. (H) Cell proportion of 4 state cells in young, middle age, old, geriatric, control (12month) and TPCA-1 (12 month) groups. (I) Cell density distribution of CD49f^high^EpCAM^low^ cells in control (12month) and TPCA-1 (12month) groups along with pseudotime.

**Figure S12.**
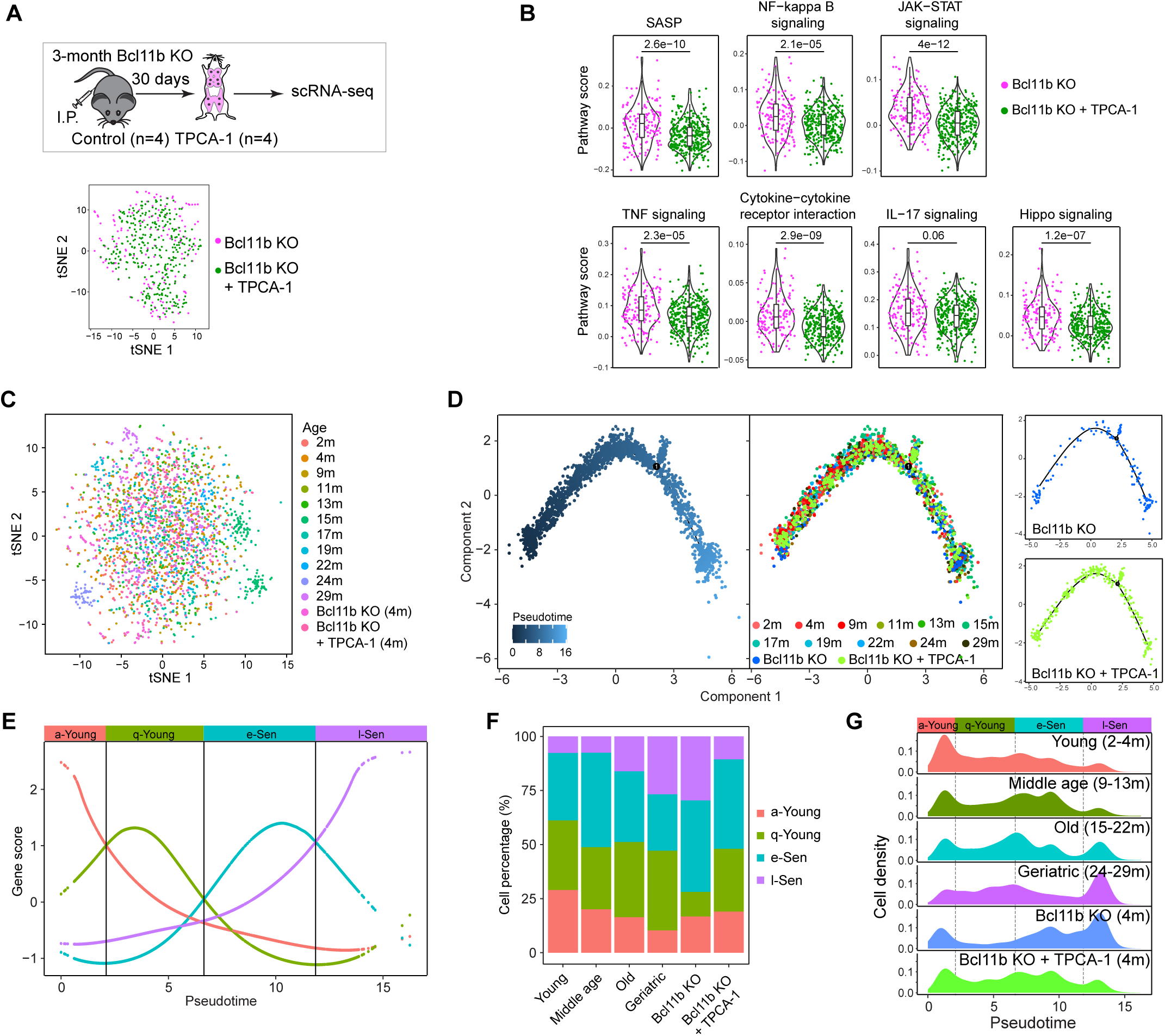
TPCA-1 can rescue Bcl11b KO derived ageing clock. (A) Schematic diagram showing the strategy of TPCA-1 (10mg/kg) treatment in vivo on 3-month-old *K14*-*Cre Bcl11b*^fl/fl^ mice. tSNE plot showing the clustering of CD49f^high^EpCAM^low^ cells from 4-month-old *K14*-*Cre Bcl11b*^fl/fl^ mice with or without TPCA-1 treatment. (B) Down-regulated signaling pathways upon TPCA-1 treatment. (C) tSNE plot showing the clustering of CD49f^high^EpCAM^low^ cells from different age WT mice and 4-month-old *K14*-*Cre Bcl11b*^fl/fl^ mice with or without TPCA-1 treatment. (D) Pseudotime-ordering analysis of single cell transcriptomes from WT mice of various ages and 4-month-old *K14*-*Cre Bcl11b*^fl/fl^ mice with or without TPCA-1 treatment. (E) Cell state signature gene scores over the pseudotime. (F) Cell proportion of 4 state cells in young, middle age, old, geriatric, control (4 month) and TPCA-1 (4 month) groups. (G) Cell density distribution of CD49f^high^EpCAM^low^ cells in control (4 month) and TPCA-1 (4 month) groups along with pseudotime.

## METHODS

### Mice and in vivo models

The Bcl11b^flox/flox^ mice (C57BL/6 background) were generously provided by Mark Leid’s lab and the B6N.Cg-Tg (KRT14-cre)1Amc/J (stock number 018964) were purchased from Jackson Laboratory. mTmG mice (B6.129(Cg)-Gt(ROSA)26Sortm4(ACTB-tdTomato,-EGFP)Luo/J, 007676) were purchased from The Jackson Laboratory (The Jackson Laboratory, Bar Harbor, Maine, USA). Female C57BL/6 mice, 2-6 months old were purchased from Jackson Laboratory, and were maintained till 29 months.

Animals were housed in a specific pathogen-free conditions and fed standard mouse chow. All animal experiments were carried out in compliance with China laws and regulations. The local institutional animal ethics board approved all mouse experiments (permission numbers: 19-001-2-CS). Experiments were performed in accordance with government and institutional guidelines and regulations.

### Tissue Processing and Flow Cytometry

Mammary glands were collected from 2nd, 3rd, 4th pair of mammary glands of C57/BL6 mice, and were dissected and processed according to the published protocol ^80^ with minor revision. Mammary glands were minced into 1 mm^3^ size using a tissue cutting blade and digested with 0.5 mg/mL Collagenase type III (Worthington, LS004182) and 50 U/mL hyaluronidase (Worthington, LS002592) for 2 hours with gentle pipetting every 30 mins. Digested mammary homogenate was collected and treated with ACK lysing buffer for 5 mins on ice, then was digested using 0.25% Trypsin-EDTA (GIBCO) for 5 mins, followed by DNase I (Worthington, LS002139) digestion. After filtered by 70 μm strainer, the dissociated mammary cells were stained with CD45 (Biolegend), CD31 (Biolegend), Ter119 (Biolegend), EpCAM (Biolegend), CD49f (Biolegend) for 20 mins on ice. Cells were washed and resuspended in HBSS+2% FBS+1% PSA+DAPI (1 μg/mL), then were sorted using FACS Aria II (BD Bioscience). FACS data was analyzed by FlowJo (V10).

### Transplantation

For transplantation assay of young and aged mammary gland, 3-month-old and 18-month-old WT C57BL/6 mice were lineage depleted using Mammary Epithelium Enrichment Kit (Stem Cell Technologies) and resuspended in injection media (HBSS +2% FBS+1% PSA+50% Matrigel) to 50k/5μL, and serially diluted to 20k/5μL, 5k/5μL and 2k/5μL. Cells were subjected to limiting dilution transplant to the cleared fat pad of 3-week-old recipient mice (C57BL/6). Briefly, the recipient mice were anesthetized with pentobarbital sodium at a dose of 70 mg/kg. The inguinal rudimentary tree was removed and 5 μL of cell suspension was injected onto the residual fat pad using 25 μL Hamilton Syringe. 7 weeks later, the recipient mice were analyzed using mammary gland whole mount carmine staining. The MRU frequency and confidence interval were determined by ELDA (http://bioinf.wehi.edu.au/software/elda/).

For transplantation of Bcl11b knockout cells, mammary cells from 4-month-old wild-type and Krt14-cre Bcl11b^flox/flox^ mice were resuspended in injection media (HBSS+2% FBS+1% PSA+50% Matrigel) to 200k/5μL. Cells were injected into the cleared fat pad of 3-week-old recipient mice (C57BL/6). 7 weeks later, the recipient mice were subsequently mated with WT male mice for 3 rounds. Mammary Tissues were collected at designated time points and subjected to whole mount analysis.

For transplant assay of IKKb overexpression cells, CD49f^high^EpCAM^low^Lin^-^ cells from 2-month-old C57BL/6 WT mice were sorted using FACS and resuspended in culture media (DMEM/F12+2% FBS+1% PSA+2% B27) supplemented with EGF (10 ng/ mL, BD Bioscience), Rspo1(250 ng/mL, R&D), ROCK inhibitor Y27632 (10 uM, Sigma) to 100k/200μL/well in 96-well non-adherent culture plate (Corning). Cells were transduced with lentivirus-pCDH or lentivirus-pCDH-IKKb at MOI 20 overnight. The next day, cells were seeded on top of the Matrigel to do the colony culture. One week later, colonies were digested using DispaseII 1 mg/mL (Sigma, D4693) for 1 hour followed by treatment of 400 μL TrypLE™ Select (GIBCO)/eppendorf tube at 37℃ for 5mins. Dissociated cells were neutralized with HBSS+2% FBS+1% PSA and stained with DAPI. GFP+ cells were sorted and then resuspended with injection media (HBSS+2% FBS+1% PSA+50% Matrigel) to 2.5k/5μL, and serially diluted to 500 cells/5μL, 100 cells/5μL and 50 cells/5μL. Cells were subjected to limiting dilution transplant to recipient mice. Mice were maintained in aseptic sterile condition for 7 weeks before whole mount analysis. For secondary transplant, mammary fat pad at 2.5k/5μL dilution, which exclusively gave rise to full tree, were collected and digested to single cell suspension. Cells from one fat pad of 2.5k/5μL dilution group were divided equally to six parts and transplanted to 6 cleared fat pad of recipient mice, respectively. Mice were maintained for 7 weeks before whole mount analysis. The mammary gland area percentage was determined by the outgrowth area divided by cleared fat pad area. The area was measured by Image J software.

### Colony formation assay

For colony formation assay, 35μL/well growth factor reduced Matrigel (Corning, 356231) was overlaid on the 96-well plate and solidified at 37℃ for 10 mins. CD49f^high^EpCAM^low^Lin^-^ cells were collected from FACS and cultured in 200 μL culture media (DMEM/F12+2% FBS+1% PSA+2% B27) supplemented with 10 ng/ mL EGF (BD Bioscience), 250 ng/ mL Rspo1(R&D), 10 μM ROCK inhibitor Y27632 (Sigma), and then were plated on top of the Matrigel. Cells were cultured at 37℃ incubator with 5% CO_2_ for 7 days.

For colony formation ability test of aging CD49f^high^EpCAM^low^Lin^-^ cells, CD49f^high^EpCAM^low^Lin^-^ cells were sorted from 2-4 months, 12-13 months and 22-29 months mice, seeded on Matrigel in 96-well plates, 3000 cells per well, three replicates for each mouse sample. Colony number was counted after 7 days culture.

For colony formation assay of IKKb overexpression cells, mammary CD49f^high^EpCAM^low^Lin^-^ cells were sorted from 2-4 months WT or K14-Cre Bcl11b^fl/fl^ mice and transduced with pCDH and pCDH-IKKb virus for 7 days. 9000 GFP positive cells were sorted and seeded to 3000 cells/200μL/well equally in three wells in the culture medium. Colony number was counted after 7 days culture. Each group was repeated with three biological replicates.

### Mammary gland whole mount carmine staining

Mammary gland was dissected and fixed in Carnoy’s solution (60% Ethanol, 30% CHCl_3_,10% Acetic Acid) for 4hrs, washed with 70% ethanol and ddH_2_O, and then stained with carmin-alum staining solution (0.2% wt/vol carmine (Sigma, C1022), 0.5% wt/vol aluminum potassium sulfate (Sigma, A7176) and 0.01% wt/vol thymol in ddH_2_O) overnight. Tissue was dehydrated through 75%, 95% and 100% ethanol, and then placed in xylene to remove the fat tissue. For long-term preservation, tissue was mounted by Permount® (Fisher Scientific). Images were obtained using a stereomicroscope (Nikon, SMZ18).

### β-Galactosidase staining

Mammary gland was dissected and immediately fixed by 2% formalin containing 0.25% glutaraldehyde for 1.5 hours followed by 30% sucrose infiltration overnight, and then was embedded in O.C.T. compound (Tissue-Tek) and frozen at −80℃. Frozen tissue was sectioned to 14 μm using Cryostat Leica CM3050 S (LEICA). Frozen sections were hydrated with PBS for 10min at RT and stained using Senescence β-Galactosidase staining kit according to manufacturer’s instructions (Cell signaling technology, 9860). The sections were incubated at 37℃ in a dry incubator (no CO_2_) for 48 hours and photographed.

### Western Blot

Comma D beta cell line ^81^ was kindly provided by Dr. Medina, and was described previously. Comma D beta cell line was cultured in DMEM-F12 (Invitrogen) supplemented with 2% of Fetal Bovine Serum (Hyclone), 1% PSA (Invitrogen), 10 ng/mL EGF(BD) and 5 μg/mL Insulin (Sigma), at 37℃ with 5% CO2. Comma D beta cells were harvested and lysed using 2X Laemmli SDS sample buffer (100 mM Tris pH6.8, 10% glycerol, 4% SDS, 0.01% Bromophenol Blue), and boiled on heat block at 100℃ for 15 mins. Samples were loaded to 4%–20% precast gradient gel (Bio-Rad) and electrophoresed at 200v for 45min, and transferred to Odyssey® nitrocellulose membrane (LI-COR). After being blocked by PBS+0.1% tween 20+5% Non-fat dry milk for 1hour at RT, the membrane was subjected to primary antibody staining beta Actin (Santa Cruz), Rat anti-Bcl11b (abcam, ab18465), Rb anti-IKKβ (Cell signaling technology, 2370), Rb anti-p-IKKα/β (Cell signaling technology, 2697), Rb anti-p-p65 (Cell signaling technology, 3033), Rb anti-p65 (Cell signaling technology, 8242), Mouse anti-IκBα (Cell signaling technology, 9247) at 4℃ overnight. Membrane was washed using PBST (PBS+0.1% Tween 20) 3×10min, and stained with secondary antibodies HRP-Donkey anti mouse (Cell signaling technology, 7076S), HRP-Donkey anti rat (Cell signaling technology, 7077S) or rabbit (Cell signaling technology, 7074S) 1:10,000 at RT for 1 hr. Membrane was subsequently washed 3×10 mins by PBST and developed using SuperSignal® West Dura Extended Duration Substrate (Thermo Scientific, 34094) and imaged by Gel imaging system (GE, AI680RGB).

### RNA extraction and Real-Time PCR

For the TNFα and LPS treatment assay, after starvation for 12 h, pInducer-Bcl11b Comma D beta cells were treated with 50 ng/mL doxycycline for 12h to induce Bcl11b expression. Cells were then treated 20 ng/mL TNFα or 10^3^ EU/mL LPS-EB for 10 hours before lysed by 400 μL Trizol (Life Technologies),. RNA was extracted according to the manufacturer’s instruction with addition of ultrapure glycogen (Thermo Scientific, R0551) as carrier. RNA was reverse transcribed to cDNA using PrimeScript^TM^ RT reagent kit (TaKaRa, RR037A) according to the manufacturer’s instructions. cDNA was then subjected to the real time PCR for specific gene target by TB Green Premix Ex Taq (TaKaRa, RR420B) according to manufacturer’s instructions using Real Time PCR system (SIS-PCR005, Jena).

For the drug screening assay, Comma D beta cells were treated with Rapamycin (0.1 μM, 1 μM, 10 μM, Selleck, S1039), Dactolisib (0.1 μM, 1 μM, 10 μM, Selleck, S1009), LY2409881 (0.1 μM, 1 μM, 10 μM, Selleck, S7697), BMS-345541 (0.01 μM, 0.1 μM, 1 μM, Selleck, S8044), QNZ (0.01 μM, 0.1 μM, 1 μM, Selleck, S4902), Asprin (1 μM, 10 μM, 100 μM, Selleck, S3017), C188-9 (0.1 μM, 1 μM, 5 μM, Selleck, S8605), SH-4-54 (0.01 μM, 0.1 μM, 1 μM, Selleck, S7337), TPCA-1 (1 μM, 5 μM, 25 μM, Selleck, 2824), Scutellarin (1 μM, 5 μM, 25 μM, Selleck, S3810), IWR-1-endo (0.1 μM, 1 μM, 5 μM, Selleck, S7086), LY411575 (0.1 μM, 1 μM, 10 μM, Selleck, S2714), Vismodegib (0.1 μM, 1 μM, 5 μM, Selleck, S1082) and Sonidegib (0.1 μM, 1 μM, 10 μM, Selleck, S2151) for 24 hours. For primary mammary cell validation experiment, CD49f^high^EpCAM^low^Lin-cells from 4-month-old, 14-month-old, 24-month-old WT mice and 4-month-old K14-Cre Bcl11b^fl/fl^ mice were sorted into the 96-well Ultra-Low attachment culture plate (Corning, 3474) (10,000 cells/200μL/well) and cultured in the culture media (DMEM/F12+2% FBS+1% PSA+2% B27+10 ng/mL EGF+250 ng/ mL Rspo1+10 μM Y27632) with 25 μM TPCA-1 or DMSO for 24 hours. DAPI negative cells were then sorted to perform RT-PCR assay. Data was analyzed by Excel and GraphPad Prism 7.00. The relative gene expression was normalized by β-actin expression. Target gene Sybrgreen primers were designed by IDT (Integrated DNA Technologies) and the primers are listed below: Bcl11b (Exon1-2): Forward ATGCCAGAATAGATGCCGG, Reverse CTCTATCTCCAGACCCTCGTC; Bcl11b (Exon2-4): Forward AGGAGAGTATCTGAGCCAGTG, Reverse GTTGTGCAAATGTAGCTGGAAG; Irak2: Forward TGTCACCTGGAACTCTACCG, Reverse TTTCTCCTGTTCATCCTTGAGG; Tnfrsf1b: Forward ACTCCAAGCATCCTTACATCG, Reverse TTCACCAGTCCTAACATCAGC; β-actin: Forward ACCTTCTACAATGAGCTGCG, Reverse CTGGATGGCTACGTACATGG.

### Luciferase assay

Comma D beta cell line which was stably expressed *pInducer Bcl11b* plasmid and NFκB-inducible Luciferase reporter plasmid was used for this experiment. After starvation for 12 hours, cells were treated with Doxycycline (Sigma, D9891) at various doses followed by 20 ng/mL TNFα (Biolegend, 575204), 1 μg/mL LPS-PG (Invivogen, tlrl-ppglps), 10^3^ EU/mL LPS-EB (Invivogen, tlrl-3pelps), and 100 ng/mL PMA (InvivoGen, tlrl-pma) treatment, as indicated. Cells were washed with 1× PBS and lysed with PLB according to the manufacturer’s instruction for 15 mins (Promega, E1910). Cells were transferred to a 96-well plate, 20 μL/well. After added with 100 μL/well LAR II, the sample was measured by the luciferase activity using the microplate reader (Thermo, Varioskan LUX).

### Immunofluorescence

For frozen section, mammary gland was dissected and immediately fixed using 4% formalin for 2 hrs followed by PBS washing and 30% sucrose infiltration overnight. The fixed mammary tissue was then embedded in O.C.T. compound (Tissue-Tek) and frozen at −80℃. Frozen tissue block was sectioned to 14 μm at −35℃ using Cryostat Leica CM3050 S (LEICA). For immunofluorescence assay, frozen sections were rehydrated with PBS for 10 mins at RT. Sections were blocked with TBS+0.1% Triton X-100+2% BSA+10% Donkey serum for 1 hr at RT, and then stained with primary antibody mouse anti-p16^Ink4a^ (Santa Cruz, sc-1661), Rb anti-p65 (Cell signaling technology, 8242), Rat anti-Bcl11b (Abcam, ab18465) 1:200 overnight at 4℃. After washed by TBST (TBS+0.1%Triton X-100) three times, sections were stained with secondary antibody Donkey anti-mouse, rat, rabbit 1:200 (Jackson ImmunoResearch) for 1 hr at RT. After 3×TBST washing and brief 1 μg/mL DAPI staining, sections were mounted with Antifade Mounting Medium (Beyotime).

### Immunohistochemistry and HE staining

Mammary tissue was collected and immediately fixed using 4% formalin overnight at 4℃ and dehydrated by gradient ethanol solution (70%, 85%, 95%, 100%). Dehydrated tissue was infiltrated by Xylene solution and embedded with paraffin. Tissue block was sectioned to 5 μm using Rotary Microtome Leica RM2255 (LEICA). Paraffin section was de-paraffinized using Xylene and rehydrated followed by gradient ethanol solution (100%, 95%, 85%, 70%, 0%) and subjected to immunohistochemistry staining according to the Histostain-Plus IHC Kit (NeoBioscience, ENS003.120, ENS004.300). Antigen was retrieved in citrate buffer (10 mM Sodium Citrate, 0.05% Tween 20, pH 6.0) for 20 mins at 100℃ in microwave. The sections were treated with 3% H_2_O_2_ for 10 mins, washed and blocked for 1 h, sections were incubated with Rb anti-p-p65 (Abcam, ab131100), Rb anti-Il-6 (NOVUS, NB600-1131), Mouse anti-Ssea1 (Abcam, ab16258) and Rb anti-Oct4 (Abcam, ab19857) overnight at 4℃. Then sections were washed using TBST, secondary antibody incubation, HPR incubation, DAB incubation, Hematoxylin dyeing, gradient dehydration and mounted by CV5030 CoverSlipper (LEICA). Images were obtained using Eclipse Ti2 inverted microscope (Nikon).

For the HE staining assay, after de-paraffinize and rehydration, paraffin section was stained using ST5020 muti-stainer (LEICA) and mounted by CV5030 CoverSlipper (LEICA).

### ChIP-seq

8×10^7^ pInducer *Bcl11b* Comma D beta cells were treated with 100ng/mL Doxycycline overnight and harvested. Rabbit anti-IgG (Abcam, ab172730) and rabbit anti-Bcl11b (Benthyl laboratories.inc, A300-384A) were used for ChIP-seq pull-down. Briefly, after cross-linked by 1% (wt/vol) formaldehyde solution, cells were quenched by glycine (0.12M), washed one time with PBS and resuspended in PBS. Cells were lysed using SDS lysis buffer (50mM Tris-HCl 8.0, 5mM EDTA 8.0, 0.1%SDS and 1× protease/phosphatase Inhibitor Cocktail (CST, 5872S)). Then chromatins were sheared with AFA Focused-ultrasonicator using Covaris ME220 with 70 peak power, 20% duty factor, 14 average power for 3 min at 1×10^7^ cells/tube. Add 9× ChIP dilution buffer (50mM Tris-HCl 8.0, 167mM NaCl, 0.11% Triton X-100, 0.11% Sodium Deoxycholate and 1× protease/phosphatase Inhibitor Cocktail) to the sonicated chromatin. 10% of the slurry was taken as input and 90% sonicated chromatin were divided equally to two parts. The sonicated chromatin was incubated in the cold room overnight with 50μL protein G-Dynabeads (Invitrogen,10004D) which had been conjugated with 50μg appropriate IgG or Bcl11b antibody. Beads were then washed with RIPA buffer 1(50mM Tris-HCl 8.0, 150mM NaCl, 1mM EDTA, 1% Triton X-100, 0.1% SDS, 0.1% Sodium Deoxycholate and 1× protease/phosphatase Inhibitor Cocktail), RIPA buffer 2 (50mM Tris-HCl 8.0, 500mM NaCl, 1mM EDTA, 1% Triton X-100, 0.1% SDS, 0.1% Sodium Deoxycholate and 1× protease/phosphatase Inhibitor Cocktail), LiCl buffer (100mM Tris-HCl 8.0, 250mM LiCl, 1mM EDTA, 0.5% NP-40, 0.5% Sodium Deoxycholate and 1× protease/phosphatase Inhibitor Cocktail) and TE buffer (10mM Tris-HCl 8.0, 1mM EDTA), and were eluted into 200uL of ChIP direct elution buffer (10mM Tris-HCl 8.0, 300mM NaCl, 5mM EDTA, 0.5% SDS, 0.2% Sodium Deoxycholate). Samples were reverse cross-linked at 65℃ overnight, treated with 4uL RNase A (1mg/mL) at 37℃ for 30min, incubated with 1uL proteinase K (10mg/mL) at 55℃ for 1h, and extracted by phenol/chloroform. ChIP DNA library was constructed by VAHTS Universal DNA Library Prep Kit for Illumina V2 (Vazyme, ND606). Briefly, samples were subjected to adapter ligation, ChIP DNA was then amplified 12 cycles and purified using AMpure XP beads (Beckman, A63881) twice and then were submitted for150 bp paired-end sequencing on an Illumina novaseq 6000 platform (Novogene).

### Single cell RNAseq library preparation and sequencing

A modified Smart-seq2 protocol was applied for single-cell RNA-seq^42,43^ according to previously reported protocol. Briefly, single CD49f^high^EpCAM^low^Lin^-^ cell from various age (2,4,9,11,13,15,17,19,22,24, 29 months old) mice or DMBA tumors were directly sorted into 96-well plate containing lysis buffer (0.05uL RNase Inhibitor (40 U/μL), 0.095uL 10% Triton X-100, 0.5uL dNTP (10mM), 0.1uL ERCC (3×10^5^) and 0.555 uLNuclease-free water) using FACS. Single cell was immediately lysed at 70℃ for 3 mins in the PCR system (TAdvanced 96SG, analytikjena). The sample was reverse transcribed to cDNA using SuperScriptIIreverse transcriptase (invitrogen, 18064-071) with a template switch oligo (TSO) primer and a sample-specific 25 nt oligo dT reverse transcription primer (TCAGACGTGTGCTCTTCCGATCTXXXXXXXX-NNNNNNNN-T25, X representing sample-specific barcode and N representing unique molecular identifier (UMI)). Then the cDNA was amplified by 18 cycles of PCR with 3’P2 primer and IS primer using KAPA HiFi HotStart Ready Mix (Kapa Biosystem, KK2602). After being pooled together and purified by AMpure XP beads (Beckman, A63881) twice, the barcoded DNAs were amplified using biotinylated pre-index primers by 4 cycles of PCR to introduce biotin tags to the 3′ ends of the amplified cDNAs. After purified by AMpure XP beads, cDNA was sonicated to approximately 300 bp fragments using Covaris ME220. 3’ terminal of the cDNA was enriched using Dynabeads® MyOne Streptavidin C1 beads (Invitrogen, 65001). The RNAseq libraries were constructed using the Kapa Hyper Prep Kit (Kapa Biosystem, KK8504) according to the manufacturer’s instructions. Briefly, after end repair and A-tailing, Streptavidin conjugated DNA was ligated to the appropriate concentration adapter (1:10) (NEB, 7335L). Then, after USER enzyme treatment and post-ligation cleanup, DNA was amplified with QP2 primer and short universal primer by 6 cycles of PCR and released from the streptavidin beads. Finally, AMpure XP beads were used to purify the DNA and DNA library quality was verified by Fragment Analyzer-12/96 (AATI) and then the DNA library was submitted to 150 bp paired-end sequencing on an Illumina NovaSeq 6000 platform (Novogene).

### DMBA induced tumor formation

Mouse mammary tumors induced by MPA and DMBA assay were performed according to the previous published paper ^52,53^ with minor modifications. Briefly, the cleared fat pad of 3-week-old female mice (C57BL/6) were transplanted with WT or K14-cre Bcl11b^fl/fl^ cells according to the transplantation assay methods described above in the Transplantation session. The recipient mice were implanted subcutaneously with a 50 mg 90-day-release MPA pellet (Innovative Research of America, NP-161-50mg). Three weeks later, DMBA (200μL, 5 mg/mL) (Sigma-Aldrich, D3254-1G) was administered by oral gavage 4 times throughout the following 5 weeks at -4, -3, -1, 0 weeks. Tumors were determined by manual palpation. Cancer incidence was calculated by the number of tumor cases within a designated latency period. The latency time was calculated from the last DMBA treatment day.

To test TPCA-1’s (GW683965) (Selleck, S2824) ^72,73^ effect on mammary ageing, 12-month-old virgin female WT mice or 3-month-old K14-cre Bcl11b^fl/fl^ mice were treated by 10 mg/kg TPCA-1 or 4% DMSO in PBS intraperitoneally every day for 30 days. Mammary cells were then dissociated and harvested for single cell RNAseq and pseudotime analysis.

To test TPCA-1’s effect on DMBA induced tumor formation, female mice induced by MPA and DMBA, then followed by treatments with 10 mg/kg TPCA-1 or 4% DMSO in PBS intraperitoneally every day for 7 weeks. Tumors were determined by manual palpation. Cancer incidence was calculated by the number of tumor cases within a designated latency period. The latency time was calculated starting from the last DMBA treatment day.

To test the vulnerability of ageing mammary cells to cancer, we transplanted 3 month old young mixed with old mammary cells to syngeneic recipient mice. Briefly, to avoid the influence of the environmental stromal cells, we transplanted 2 month old young mammary cells (wt) with 24 month old mammary cells (with GFP reporter) to syngeneic recipient mice at equal MRUs and induced cancer formation by DMBA treatment. We repeat this experiment with 2 month old young mammary cells (with tdTomato reporter) and 12 month old mammary cells (wt).

To test whether basal cells could be the origin of DMBA induced cancer using Krt14rtTA-TetOcre-mTmG mice. We first labeled basal cells with doxycycline induction, and then treat mice with DMBA.

### Single cell RNA-seq analysis

Raw reads were first processed using TrimGalore (https://github.com/FelixKrueger/TrimGalore) to remove adaptor sequences with paired end mode and default parameter. Quality control was evaluated with FastQC(https://www.bioinformatics.babraham.ac.uk/projects/fastqc/). For each sequencing batch, the top 100 barcode ranked by read counts were retained using whitelist tool in UMI-tools ^82^. Then barcode and UMI information were extracted from read2 using extract tool in UMI-tools and added to read1. Subsequently, read1 were aligned to the mm10 genome using STAR aligner with default parameter except for the outFilterMultimapNmax=1. Mapped reads were assigned to genes using featureCounts ^82^. Finally, we used count tools in UMI-tools to generate count matrix, which the number of UMIs represents the transcript number of each gene within each individual cell.

After getting the count matrix, we applied four criterions to further exclude cells with low data quality: first, cells with barcode not included in the barcode sequence list were removed; second, cells with ERCC percentage larger than 10% or mitochondria percentage larger than 5% were filtered out; third, cells with gene number less than 200 or more than 6000 were removed; lastly, cells with stromal gene (Pecam1, Ptprc, Lyve1, Col1a1) expression level higher than 0.1 were removed. As for gene, we removed those detected in less than 10 cells. Finally, for young and aged single cell RNA-seq assay, 1981 cells passed the quality control. The same filtering criterion was used for K14-Cre Bcl11b^fl/fl^, tumor and TPCA-1 treatment single cell RNA-seq assay. 149 K14-Cre Bcl11b^fl/fl^ CD49f^high^EpCAM^low^Lin^-^cells, 310 CD49f^high^EpCAM^low^Lin^-^ tumor cells, 373 DMSO treated and 367 TPCA-1 treated CD49f^high^EpCAM^low^Lin^-^ cells were used for analysis.

We used the Seurat v4 ^83^ to carry out the abovementioned filtering, data normalization, and all downstream analysis including dimensionality reduction, clustering, tSNE plot overlaying, and differential gene expression. More specifically, UMI counts in each cell were normalized with NormalizeData function with default parameter and vars.to.regress parameter was used to regress out the cell cycle, ERCC percentage, mitochondria percentage, batch and gene number effect. For dimension reduction, RunPCA function was used and the top 10 principal components were passed to tSNE analysis by RunTSNE function. Finally, clustering was performed by FindNeighbors and FindClusters functions with original Louvain algorithm and the resolution was set to 1.

### Single cell trajectory analysis

Single cell trajectory was inferred by Monocle2 ^84^. Count matrix was input to Monocle2 and negative binomial distribution was used for building statistical distribution for read counts with lower detection limit set as 0.5. PhenoData was exported from Seurat object. Dimension reduction was done with DDRTree and the effect of Size_Factor, num_genes_expressed, ERCC percentage, mitochondria percentage, cell cycle and batcheffect were regressed out. The root of the pseudo-time trajectory was selected as the most abundant cell state in the cells from the 2-month-old batch. Genes significantly changed along pseudo-time were selected using differentialGeneTest function with qval<0.05, resulting in 1932 differentially expressed genes. A new count matrix was generated from the original to include only differentially expressed genes. The count matrix was then smoothed using genSmoothCurves function, log transformed using a base of 10 and pseudo-count of 1 to prevent logarithm of zero value. All elements in the transformed count matrix were further truncated by a straightforward way that all elements larger than 3 or smaller than -3 were set as 3 or -3 respectively. Gene hierarchical clustering was performed on the transformed matrix with Heatmap in ComplexHeatmap package ^85^. Genes were clustered into 4 clusters according to gene expression in cells along with pseudo-time increasing. Based on intersecting point of the average expression of genes in each cluster, cells were separated into 4 states. The same pipeline for combined aging & Bcl11b ko data and aging & TPCA-1 treatment data.

### Pathway enrichment analysis

All gene symbols were mapped to their Entrez gene ids using biomaRt ^86,87^. Then, both GO and KEGG pathway enrichment analysis were performed by clusterProfiler ^88^. For simplicity, only biological process (BP) terms in GO were used for enrichment analysis. To do GSEA analysis, genes ordered according to decreased foldchange were fed to GSEA function. To do GSEA enrichment of Bcl11b-KO vs WT tumor CD49f^high^EpCAM^low^Lin^-^ cells on 4 state marker genes, marker genes of each state were considered as one pathway. For enriched results from enrichGO, enrichKEGG or GSEA, only pathways with BH-adjusted p value less than 0.05 were retained. Pathway activity of each cell was got with AddmoduleScore function in Seurat package ^83^. When showing pathway activity along with pseudotime, the activity was also smoothed with genSmoothCurves function in monocle package ^84^.

### Cell cycle state determination

We re-implemented the method previously used in Kowalczyk et al ^46^ to determine the cell cycle state for each cell. First, filtered count matrix after Seurat was transferred into TPM using calculateTPM function in scater package ^89^ and then log-transformed by log_2_(TPM+1). The cycle gene list in human was taken from the previously published paper by Whitfield ^90^. All human gene symbols were transformed to mouse using biomaRt package ^86,87^. Genes were filtered by the correlation with the average gene expression of corresponding cell cycle stage. To retain mammary gland specific cell cycle genes, the correlation threshold was set to 0.25. Finally, we got 14, 13, 21, 19, 13 genes for G1/S, G2, G2/M, M/G1 and S phase, respectively. The average cell cycle gene expression of each stage was calculated. We identified cells with G1/S score <0 & G2/M <0 as G0 cells. Other cells were defined as the corresponding stage according to gene maximum expression.

### SASP gene score calculation

The SASP gene list in human was taken from the previously published paper by Coppe et al ^11^. All human gene symbols were mapped to mouse symbols by biomaRt package ^86,87^. Then, the SASP score for each cell were calculated using AddModuleScore function in Seurat package ^83^.

### Pathway score and transcription factor activity score calculation in human BRCA

BRCA fpkm data was downloaded with TCGAbiolinks ^91^, in which we only used the paired samples, then pathway score was calculated as mean expression of pathway genes. The result was shown with ggplot2 (https://ggplot2.tidyverse.org). Statistical analysis was performed using two-tailed paired t-tests. To calculate transcription factor activity, we used the transcription factor target genes from Transcription Factor Target Gene Database ^92^.

### scEntropy analysis

We compute single cell entropy according to the public paper ^55,93^. Briefly, single cell count expression profile from R was exported as .mat format which can be loaded into MATLAB. For calculating the entropy of cells from different ages, we constructed the gene co-expression network and apply it to Bcl11b ko cells. After computing the entropy of each cell, we visualize these results in R with ggplot2.

### Transcription factor activity analysis

We got transcription factor target genes from the following three public databases: ENCODE ^94^, ChEA ^95^, and TRRUST v2 ^96^. For each cluster of genes changing along with pseudo-time, TFs were enriched with hypergeometric test which was implemented with phyper function in R. Transcription factor activity was calculated with transcription factor target genes from the combined databases using AddModuleScore function in Seurat package ^83^. To calculate Bcl11b activity score, we did ChIP-seq assay for Bcl11b to find Bcl11b target genes. Based on scRNA-seq of wild type CD49f^high^EpCAM^low^Lin^-^ cells and Bcl11b ko CD49f^high^EpCAM^low^Lin^-^ cells from 4-month-old mice, we got Bcl11b positively regulated genes (genes with pval<0.05, ko < wt intersect with Bcl11b targets) and negatively regulated genes (genes with pval<0.05, ko > wt intersect with Bcl11b targets). Then Bcl11b activity was computed as weighted average expression of the negatively regulated genes.

### ChIP-seq data processing

ChIP-seq reads were trimmed with TrimGalore(https://github.com/FelixKrueger/TrimGalore) with paired end mode and default parameter and QC was done with FastQC(https://www.bioinformatics.babraham.ac.uk/projects/fastqc/). Trimmed reads were aligned to the mm10 mouse genome with bowtie2 (https://github.com/BenLangmead/bowtie2) with default parameter. Then, the alignment was filtered using samtools ^97^ with the following parameter: -F 1804 -f 2 -q 30. Duplicated reads were marked with MarkDuplicates in picard-tools (http://broadinstitute.github.io/picard/) and filtered out with samtools. Finally, we found bcl11b binding sites on the genome based on input signal by MACS2 using following parameter: -f BAMPE -g mm --keep-dup all --nomodel. Bcl11b binding sites were analyzed using ChIPseeker package ^98^. ChIP-seq signals were visualized using Integrative Genomics Viewer (IGV) software^99^.

### Quantification and statistical analysis

Log-rank test was used between WT and K14-Cre Bcl11b^fl/fl^ group tumor formation kinetics by GraphPad Prism 7.00. For limiting dilution analyses, the frequency of mammary repopulating unit was calculated using ELDA software ^100^. Statistical analyses were performed using GraphPad Prism 7.00 with unpaired or paired two-tailed Student’s t-test, as indicated in the figure legends. Bar graphs represent mean ± SD or mean ± SEM, as indicated.

## Acknowledgments

This work was supported by National Natural Science Foundation of China NSFC grant 32170803, 81872405. This work was supported by Westlake Education Foundation. We thank Yajing Guo for the management of mice. We thank Peng Guo, Dandan Liao and Yanan Tang at the Flow Cytometry Facility of Westlake University for assistance in FACS sorting. We thank Haiyan Zhang at the Genomics Core Facility of Westlake University for assistance in library quality test. We thank the Westlake Animal Facility for mouse husbandry. We are grateful for Dr. Dangsheng Li for valuable discussions and suggestions.

## Authors’ Contributions

S.C. and H.B. conceptualized the project and developed the methodology; H.B. performed most of the experiments; S.C. and H.B. conceptualized the bioinformatic analysis. X.L. and N.L. performed scRNA-seq and ChIP-seq data analysis; M.L. performed the immunofluorescence assay and Y.M. performed the immunohistochemical analysis.

